# S9 Protease WprP Catalyzes Uniform and Sequential Cleavage on the Precursor Peptide in RiPP Biosynthesis

**DOI:** 10.1101/2025.10.18.683210

**Authors:** Jabal Rahmat Haedar, Abujunaid Habib Khan, Stefano Donadio, Chin-Soon Phan

**Affiliations:** Latvian Institute of Organic Synthesis, Aizkraukles Street 21, LV-1006 Riga, Latvia; NAICONS Srl, 20139 Milan, Italy

## Abstract

Serine proteases in ribosomally synthesized and post-translationally modified peptides (RiPPs) catalyze the cleavage on the precursor peptides in the biosynthesis of RiPP natural products. Here, we identified an uncharacterized serine protease WprP_2_ from *Streptomyces venezuelae* NPDC049867, encoded next to the radical SAM enzyme WprB_2_ involved in the biosynthesis of cyclophane natural products. *In vitro* characterization of S9 protease WprP_2_ revealed that the precursor peptide WprA_2_ is uniformly and sequentially cleaved. The cleavage activity of WprP_2_ has not been seen in any serine proteases and expands the S9 protease in RiPP biosynthesis.

---

Ribosomally synthesized and post-translationally modified peptides (RiPPs) are a rapidly growing class of peptide natural products defined by their post-translational modifications.^1,2^ The backbone structure of RiPPs is encoded in the core peptide region of the precursor peptide, which is modified by post-translational modification enzymes (maturases) and then removed from the leader peptide by proteases and released as mature natural products (Figure 1A).^1,2^ In general, the removal of the leader peptide in RiPPs is crucial for the generation of bioactive natural products.^3^ The class I lanthipeptide nisin A is the oldest and most extensively studied RiPP natural product; isolated in 1928, with its structure determined in 1971, assigned to RiPP in 1988, it possess antimicrobial activities against Gram-positive bacteria.^4-7^ Cleavage of the leader peptide is necessary for the production of biologically active nisin.^8^

**Figure 1.**
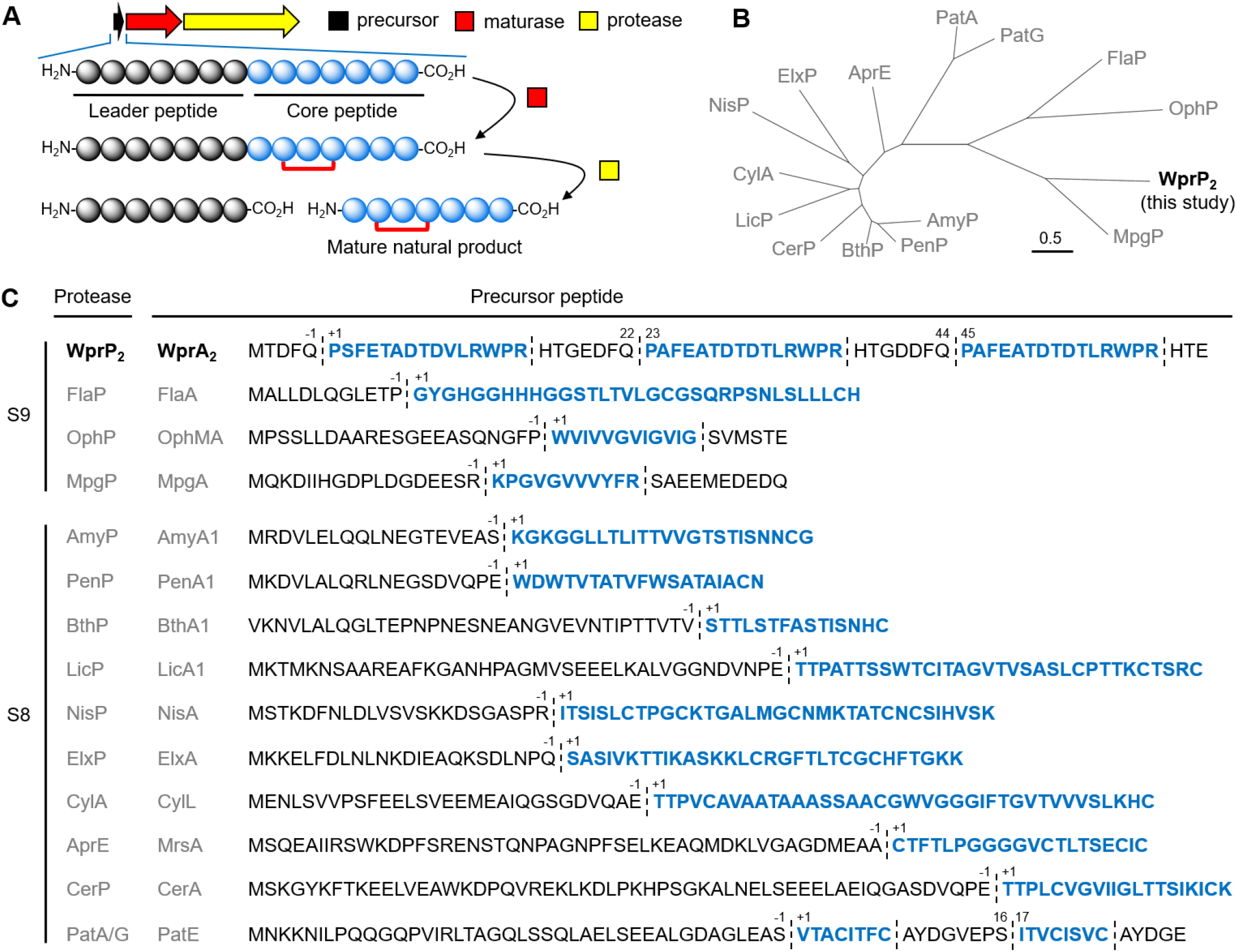
(A) Overview of RiPP biosynthesis. (B) Phylogenetic tree of known and newly characterized serine proteases in RiPP. (C) Cleavage sites of known and newly characterized serine proteases in RiPP biosynthesis. The known and newly characterized serine proteases in RiPP are shown as grey colored letters and black colored bold letters, respectively. Protease cleavage sites are shown as dashed lines. Core peptides are shown as blue colored bold letters. Residues within the precursor sequence are numbered ‘+1’ from the start of the core peptide.

Proteases involved in RiPP biosynthesis include: (1) cysteine family proteases that usually cleave after a typical double glycine-like motif, (2) serine family proteases that have serine as the nucleophilic residue in the active site, (3) metalloproteases that use metal ions to catalyze cleavage, and (4) other uncategorized proteases.^9^ Over the past decade, the number of characterized serine proteases in RiPP biosynthesis has increased (Figure 1B). To date, only two groups of serine proteases involved in the maturation of RiPP natural products: (1) the S8 family which includes AmyP,^10^ PenP,^10^ BthP,^10^ LicP,^11^ NisP,^12^ ElxP,^13^ CylA,^14^ AprE,^15^ and CerP,^16^ involved in the biosynthesis of class I to III lanthipeptides, and PatA/G involved in the biosynthesis of cyanobactin;^17,18^ and (2) the S9 family, which contains only FlaP,^19^ OphP,^20^ and MpgP^21^ involved in the biosynthesis of class III lathipeptides, omphalotins, and clavusporins, respectively (Figure 1C).

Cross-linking of peptides is an important chemical feature that improves the stability and biological activity of peptides.^22,23^ An important example is the natural product, darobactin A, a heptapeptide composed of two three-residue motif cyclophanes that targets the essential outer membrane protein BamA of Gram-negative bacteria.^23,24^ Recently, radical SAM (rSAM) proteins were discovered as a group of RiPP enzymes that catalyze the formation of a three-residue motif cyclophane between one aromatic amino acid and one aliphatic amino acid on the precursor peptide.^25-27^ However, in many cases, these rSAM-RiPP biosynthetic gene clusters (BGCs) do not contain proteases,^23,28-33^ and thus their mature natural products remain elusive, with the exception of the rSAM-RiPP natural products, darobactin A and dynobactin A isolated from native strains.^23,33,34^ To date, only one protease annotated as peptidase C39 has been characterized in rSAM-RiPP biosynthesis, which cleaves after a canonical double glycine motif to generate a mature natural product, xenorceptide (Figure S1).^28^

In our previous study, we discovered the rSAM cyclophane synthase WprB_1_, which catalyzes a cross-link between Trp-C5 and Arg-Cγ at a three-residue WPR motif on the precursor peptide WprA_1_ (Figure 2A).^32^ Unlike other rSAM-RiPP precursor peptides, WprA contains three repeats of the WPR motif. However, no proteases were found in the flanking region of the BGC. We are interested in searching the proteases that can cleave the precursor peptide WprA after installing the cyclophanes at WPR motifs.

**Figure 2.**
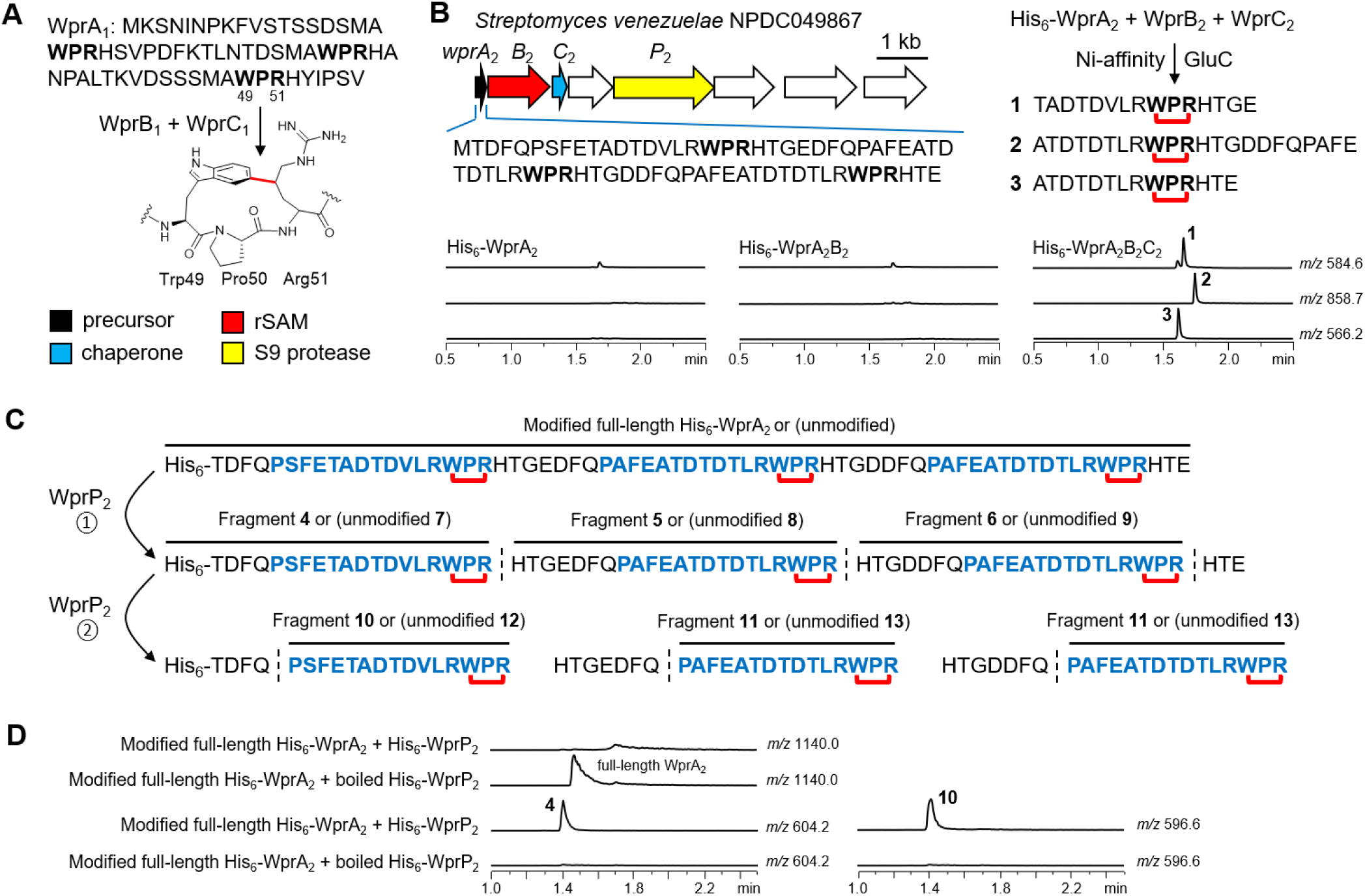
(A) Summary of previous studies of WprA_1_B_1_C_1_.^31^ (B) *In vivo* coexpression of His_6_-WprA_2_B_2_C_2_ followed by Ni-affinity purification and GluC digestion yielded peptide fragments **1**-**3**. (C) Overview of S9 protease WprP_2_ cleavage activity yielded peptide fragments **4**-**13**. (D) *In vitro* characterization of native or boiled His_6_-WprP_2_ + modified or unmodified His_6_-WprA_2_. Cross-link formation on the peptide sequences is shown as red connectors.

To this end, we used Position-Specific Iterative Basic Local Alignment Search Tool (PSI-BLAST)^35^ to expand for rSAM sequences homologous of WprB_1_.^32^ We identified 16 homologous WprB whose cognate precursor peptides contain one or three repeated WPR motifs and then searched for the presence of a protease in the flanking region (Figure S2 and Table S1). We detected a S9 protease in two distinct BGCs: one from *Streptomyces venezuelae* NPDC049867, which encoded a precursor peptide containing three WPR motifs; and the other from *Streptomyces jietaisiensis* NBC_00023, where the precursor peptide contained only one WPR motif. Based on the presence of three WPR repeats in the precursor peptide, as in our previous study, we aimed to elucidate the cleavage activity by the S9 protease WprP_2_ from *S. venezuelae* NPDC049867 (Figure 2B). WprP_2_ shares amino acid identity with FlaP (28.7%),^17^ OphP (24.3%),^18^ MpgP (31.4%),^21^ and no significant similarity with all characterized S8 proteases found in RiPP.^10-18^

We first validated the cross-linking activity at WPR motif(s) by coexpression of WprA_2_B_2_C_2_. The N-terminal His_6_ tag precursor peptide WprA_2_ expressed in *Escherichia coli* NiCo21(DE3) alone or coexpressed with rSAM enzyme WprB_2_ and/or chaperone WprC_2_, purified by Ni-affinity chromatography and digested with GluC. Comparative LC-MS analysis of the digests led to the identification of peaks **1**-**3** with −2 Da mass loss relative to the unmodified peptide fragment (Figures 2B and S3-S5). The results showed that WprB_2_ similar to WprB_1_,^32^ requires the chaperone WprC_2_ to catalyze the formation of a cross-link at all three WPR motifs on the precursor peptide. The nature of the cross-links at all three WPR motifs on WprA_2_ was assumed to be identical to those generated by WprB_1_ (43.8% amino acid identity with WprB_2_).^32^

Next, we investigate the S9 protease WprP_2_ activity through *in vitro* experiments (Figure 2C). As a substrate for biochemical characterization of WprP_2_, we used the Ni-affinity purified His_6_ tag precursor peptide WprA_2_, expressed alone or coexpressed with WprB_2_C_2_ in *E. coli* NiCo21(DE3). Separately, N-terminal His_6_ tag WprP_2_ was expressed in *E. coli* NiCo21(DE3), purified by Ni-affinity chromatography and analyzed by SDS-PAGE (Figure S6). When modified/unmodified full-length His_6_-WprA_2_ was incubated with native or boiled enzyme His_6_-WprP_2_, we observed that modified full-length His_6_-WprA_2_ was present only in the boiled enzyme but not in the native enzyme (Figures 2D and S7). The native His_6_-WprP_2_ cleaved after all three WPR motifs in both modified and unmodified substrates. This generated six peptide fragments **4**-**9** containing cross-linked or unmodified WPR motifs. Interestingly, further analysis showed that WprP_2_ can catalyze the second cleavage on peptide fragments **4**-**9** before the Pro amino acid at the 12^th^ residue preceding the WPR motif, generating shorter peptide fragments **10**-**13**, where the cleavage products of **5** (**8**) and **6** (**9**) have identical amino acid sequences (Figures 2C, 2D and S8-S13).

To understand the recognition sequence of WprP_2_, we designed two precursor variants His_6_-WprA_2_-eng1/2 with multiple mutations at D51A/T52A/D53A/T54A and D42A/F43A/P47A/E48A, and four precursor variants with a single mutation at P45A, Q44A, H38A and R59A. The results showed that the D51A/T52A/D53A/T54A and D42A/F43A/P47A/E48A mutations at the proximal WPR motif do not affect the cleavage activity by WprP_2_, but it impaired the cross-link formation by WprB_2_ (Figures S14-S15). The WprA_2_ H38A results showed that the cleavage activity of WprP_2_ does not require the His residue after the WPR motif, and thus all expected peptide fragments were detected (Figure S16). Surprisingly, a single mutation at R59A in WprA_2_ prevented cleavage at this position and generated a longer peptide fragment **19**, indicating that the first cleavage recognition sequence is the WPR-Xxx site (Figure S17). We next found that the second cleavage recognition sequence is the Gln-Pro site, as single mutation at P45A or Q44A prevented cleavage at this position and generated peptide fragments **21** (**22**) and **23** (Figures 3B and S18-S19).

**Figure 3.**
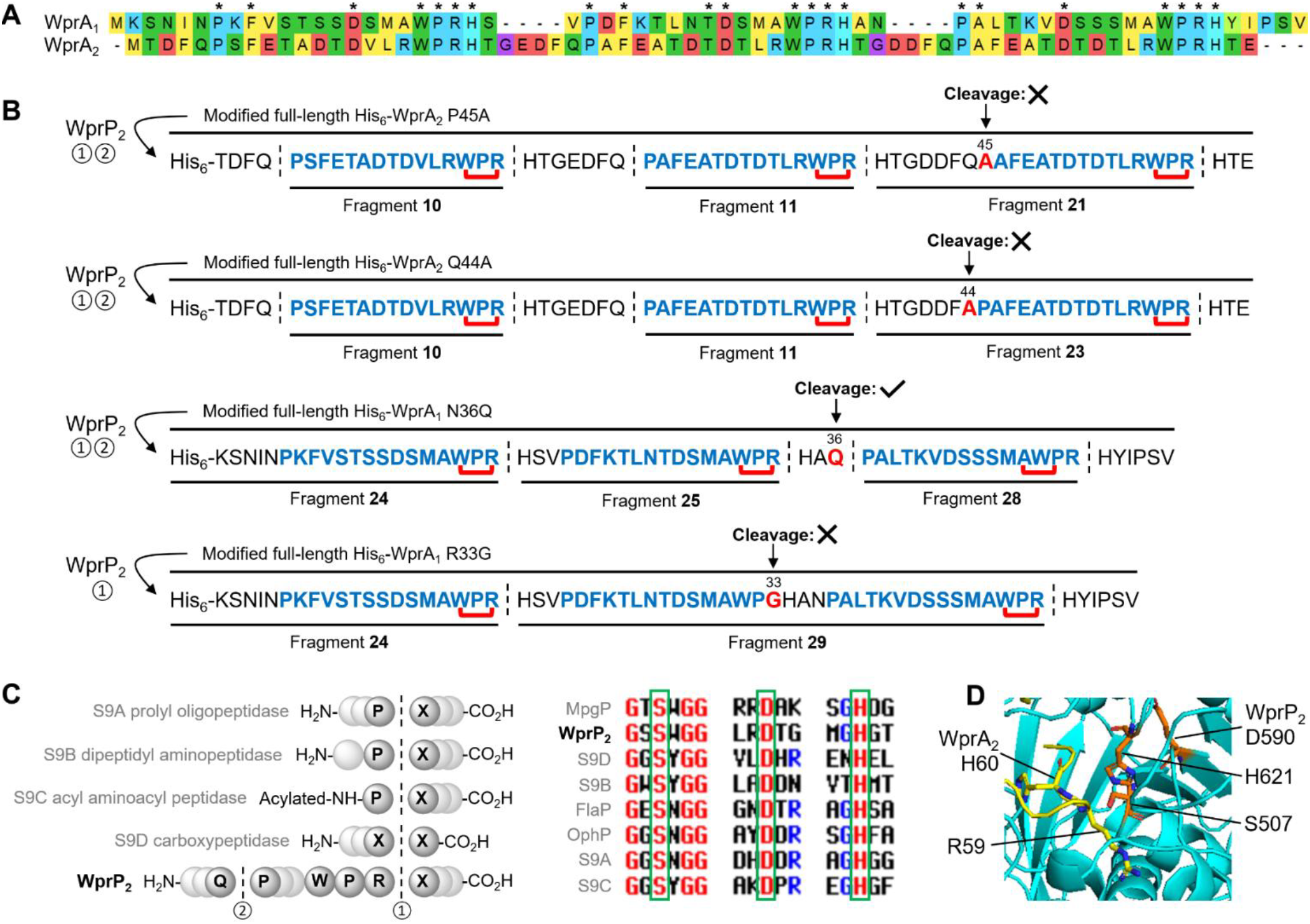
(A) Comparison of the precursor sequences of WprA_1_ and WprA_2_. (B) Recognition sequence for the cleavage activity of S9 protease WprP_2_. (C) Cleavage sites and catalytic triad of different S9 subfamily proteases. (D) AlphaFold3 predicted the complex structure of WprP_2_ with a truncated substrate of WprA_2_ showing the Arg-His cleavage site in the catalytic triad. Conserved and mutated residues on the precursor sequences are shown as asterisk symbols and red colored bold letters, respectively.

We then asked whether WprP_2_ could cleave a related substrate WprA_1_, which has different amino acid composition and length (Figure 3A). To test this, we used the modified full-length WprA_1_ in our hand for *in vitro* assay of WprP_2_. The LC-MS data detected cleavage after all three WPR motifs by WprP_2_ generating peptide fragments **24**-**26**, but we did not observe the second cleavage before the Pro residue preceding the WPR motif (Figures S20-S21). Notably, all three Pro residues in WprA_2_ are preceded by a Gln, whereas the Pro preceding amino acids in WprA_1_ are Asn, Val and Asn, respectively (Figure S22). This prompted us to design the precursor variant WprA_1_ N36Q, and as expected, a second cleavage activity was detected at the Gln-Pro site and generated peptide fragments **28** (Figures 3B and S23). To cross-validate the recognition sequence of the first cleavage, we used two precursor variants WprA_1_ with a single mutation at H52A or R33G in our hand. The results showed that WprA_1_ H52A could detect all expected peptide fragments (Figure S24). While WprA_1_ R33G prevented cleavage at this position as expected and generated a longer peptide fragment **29** containing two WPR motifs (Figures 3B and S25).

Several members of S9 family proteases are pharmacologically relevant, for example prolyl oligopeptidase is a drug target for celiac sprue and diabetes,^36,37^ while acyl aminoacyl peptidase has been reported to be associated with Alzheimer’s disease, cataract formation and cancer.^38,39^ The S9 family proteases can be classified into four subfamilies: (1) S9A prolyl oligopeptidase, which cleave peptide bonds after the Pro residue,^19,20^ (2) S9B dipeptidyl peptidase, which cleave peptide bonds at the penultimate Pro residue at the N-terminus,^40^ (3) S9C acyl aminoacyl peptidase, which cleave peptide bonds after a N-acetylated Pro residue,^41^ and (4) S9D carboxypeptidase, which sequentially cleave peptide bonds at the C-terminal residue.^42^ All S9 subfamily proteases have a signature catalytic triad consisting of Ser, Asp, and His residues (Figures 3C and S26-S27), and a single mutation at S507A, D590A or H621A completely abolished the cleavage activity of WprP_2_ (Figure S28-S29). AlphaFold3 predicted the complex structure of WprP_2_ with a truncated substrate of WprA_2_, showing that the cleavage sites of Arg-His and Gln-Pro are placed in spatial proximity to catalytic triad (Figures 3D and S30). To date, only three S9 proteases FlaP,^19^ OphP^20^ and MpgP^21^ have been characterized in RiPP biosynthesis, and WprP_2_ differs from them in three ways: (1) our phylogenetic analysis showed WprP_2_ does not form a clade with FlaP and OphP, (2) unlike known S9 proteases, WprP_2_ cleaves the peptide bond at the N-terminal of Pro residue, and (3) unlike known S9 proteases, WprP_2_ catalyzes uniform and sequential cleavage on the precursor peptide. Notably, unlike WprP_2_, commercial trypsin was unable to cleave the Arg-Xxx site of the modified full-length WprA_1_ (Figure S31), highlighting the potential applicability of WprP_2_ in cleaving peptide bond.

In conclusion, we identified an uncharacterized serine protease WprP_2_ from *Streptomyces venezuelae* NPDC049867, encoded next to the radical SAM enzyme WprB_2_ involved in the biosynthesis of cyclophane natural products. WprP_2_ catalyzes the uniform and sequential cleavage on the precursor peptide WprA_2_, with the first cleavage occurring after WPR motif and second cleavage occurring before the Pro amino acid at the 12^th^ residue preceding the WPR motif. Such cleavage has not been seen in any serine proteases from RiPP biosynthesis. Furthermore, WprP_2_ recognized both Gln-Pro and WPR-Xxx cleavage sites regardless of the amino acid composition and length in between two cleavage sites. This work unlocks a new toolkit for peptide bond cleavage and highlights its promiscuity in recognizing a wide range of substrates, providing immense potential for generating bioactive peptides.

## ASSOCIATED CONTENT

### Supporting Information

The experimental details; figures for coexpression of WprA_2_B_2_C_2_ and *in vitro* assay of WprP_2_; list of 16 homologous BGCs of *wpr*; gene sequences and primers used in this study.

## AUTHOR INFORMATION

## Author

Jabal Rahmat Haedar - Latvian Institute of Organic Synthesis, Aizkraukles Street 21, LV-1006 Riga, Latvia.

Abujunaid Habib Khan - Latvian Institute of Organic Synthesis, Aizkraukles Street 21, LV-1006 Riga, Latvia.

Stefano Donadio - Latvian Institute of Organic Synthesis, Aizkraukles Street 21, LV-1006 Riga, Latvia. NAICONS Srl, 20139 Milan, Italy.

## Notes

The authors declare no competing financial interest.

## ACKNOWLEDGMENT

This work was funded by EU project No. 101087181 (Natural Products Research at Latvian Institute of Organic Synthesis as a Driver for Excellence in Innovation). We acknowledge the supports from the LC-MS and NMR facilities, and biotechnology laboratories at Latvian Institute of Organic Synthesis.

---

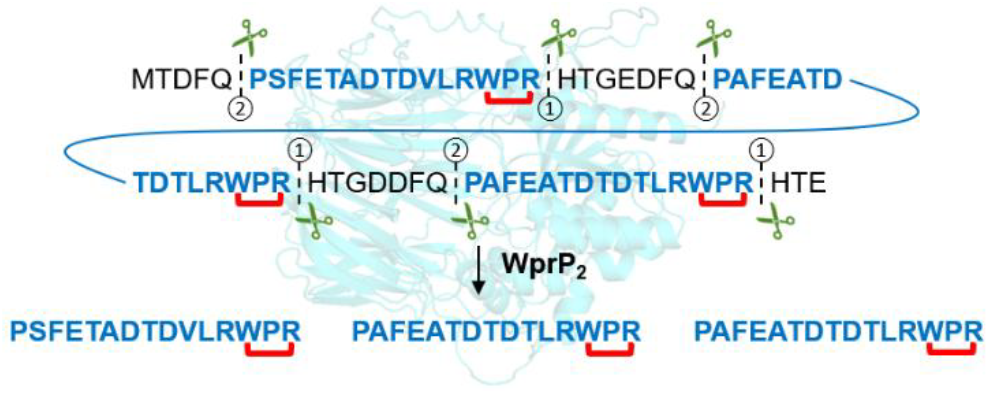

## Notes

### Competing Interest Statement

The authors have declared no competing interest.

