## Supplementary Information for "S9 Protease WprP Catalyzes Uniform and Sequential Cleavage on the Precursor Peptide in RiPP Biosynthesis"

### Table of Contents


| | |
|---|---|
| Experimental section | |
| Figure S1 | Proteases involved in the rSAM-RiPP biosynthesis |
| Figure S2 | BGCs of 16 homologous WprB |
| Figure S3 | *In vivo* coexpression of $WprA_2$ |
| Figure S4 | *In vivo* coexpression of $WprA_2$ + $WprB_2$ |
| Figure S5 | *In vivo* coexpression of $WprA_2$ + $WprB_2$ + $WprC_2$ |
| Figure S6 | SDS-PAGE of $WprP_2$ |
| Figure S7 | *In vitro* assay of $His_6$-$WprP_2$ + modified $His_6$-$WprA_2$ |
| Figure S8 | *In vitro* assay of $His_6$-$WprP_2$ + modified $His_6$-$WprA_2$ |
| Figure S9 | The MS/MS spectra of fragments **4-6** |
| Figure S10 | The MS/MS spectra of fragments **10** and **11** |
| Figure S11 | *In vitro* assay of $His_6$-$WprP_2$ + unmodified $His_6$-$WprA_2$ |
| Figure S12 | The MS/MS spectra of fragments **7-9** |
| Figure S13 | The MS/MS spectra of fragments **12** and **13** |
| Figure S14 | *In vitro* assay of $His_6$-$WprP_2$ + modified $His_6$-$WprA_2$-eng1 |
| Figure S15 | *In vitro* assay of $His_6$-$WprP_2$ + modified $His_6$-$WprA_2$-eng2 |
| Figure S16 | *In vitro* assay of $His_6$-$WprP_2$ + modified $His_6$-$WprA_2$ H38A |
| Figure S17 | *In vitro* assay of $His_6$-$WprP_2$ + modified $His_6$-$WprA_2$ R59A |
| Figure S18 | *In vitro* assay of $His_6$-$WprP_2$ + modified $His_6$-$WprA_2$ P45A |
| Figure S19 | *In vitro* assay of $His_6$-$WprP_2$ + modified $His_6$-$WprA_2$ Q44A |
| Figure S20 | *In vitro* assay of $His_6$-$WprP_2$ + modified $His_6$-$WprA_1$ |
| Figure S21 | The MS/MS spectra of fragments **24-26** |
| Figure S22 | Sequence alignment of precursor peptide WprA |
| Figure S23 | *In vitro* assay of $His_6$-$WprP_2$ + modified $His_6$-$WprA_1$ N36Q |
| Figure S24 | *In vitro* assay of $His_6$-$WprP_2$ + modified $His_6$-$WprA_1$ H52A |
| Figure S25 | *In vitro* assay of $His_6$-$WprP_2$ + modified $His_6$-$WprA_1$ R33G |
| Figure S26 | Sequence alignment of S9 proteases |
| Figure S27 | Overview of catalytic triad in S9 proteases |
| Figure S28 | SDS-PAGE of $WprP_2$ mutants |
| Figure S29 | *In vitro* assay of $His_6$-$WprP_2$ mutants + modified $His_6$-$WprA_2$ |
| Figure S30 | Predicted complex structure of $WprP_2$ with truncated $WprA_2$ |
| Figure S31 | Commercial trypsin vs $WprP_2$ against modified $His_6$-$WprA_1$ |
| Table S1 | List of 16 homologous WprB (precursor peptides/S9 proteases) |
| Table S2 | Gene sequences used in this study. |
| Table S3 | Amino acid sequence of proteins used in this study. |
| Table S4 | Primers used in this study. |
| Table S5 | Plasmids constructed in this study. |
| Table S6 | Strains used in this study. |
| Table S7 | Precursor peptides used in this study. |
| Supplementary references | |

#### EXPERIMENTAL SECTION

**General.** Chemicals and reagents were purchased from either Merck or Sigma-Aldrich unless otherwise specified. Synthetic genes inserted into expression vectors were synthesized by the Twist Bioscience (https://www.twistbioscience.com/). Gene sequences used in this study are listed in Table S2. Trypsin protease was purchased from Merck. Antibiotics (kanamycin, spectinomycin and chloramphenicol) were purchased from GoldBio. *Escherichia coli* NiCo21(DE3) and DH5α strains purchased from NEB were used for protein expression and plasmid preparation, respectively. Electroporation was carried out using a Bio-Rad MicroPulser. Terrific broth (TB) was purchased from Formedium. Luria broth (LB), Miller broth and agar were purchased from Carl Roth. SDS-PAGE, 4-20% polyacrylamide gel was purchased from GenScript. Protein marker, broad multi color pre-stained protein standard for SDS-PAGE was purchased from GenScript. The *E. coli* cells were lysed using a Hielscher Ultrasonics Sonotrode S26D7 Titanium 7mm. LC-MS experiments were performed on a Waters Acquity H-class UPLC System coupled to Waters QToF SYNAPT G2-Si Mass Spectrometer.

**Bioinformatic expansion of characterized radical SAM enzyme $WprB_1$.** To retrieve putative radical SAM enzyme sequences, Position-Specific Iterative Basic Local Alignment Search Tool (PSI-BLAST)[1] was performed in February 2025 on a non-redundant protein database in National Center for Biotechnology Information (NCBI) online database using our characterized radical SAM enzyme $WprB_1$ (WP_319939623.1)[2] as query and 1E-5 as cutoff value for two rounds of search (iterations). The putative precursor peptides were manually identified from the upstream regions of WprB. This resulted 16 homologous WprB enzymes linked to putative precursor peptides (Table S1). Further analysis of the flanking regions of these 16 homologous WprB enzymes found that two BGCs encoded S9 proteases (Figure S2), four newly identified genes were the subject of this study, these genes are designated as precursor peptide $WprA_2$ (WP_399537669.1), radical SAM enzyme $WprB_2$ (WP_399537670.1), chaperone $WprC_2$ (WP_399537671.1), and S9 protease $WprP_2$ (WP_399537673.1) as shown in Table S2.

**Phylogenetic tree construction**. The 13 characterized serine proteases, AmyP (WP_013351403.1),[3] PenP (WP_430242725.1),[3] BthP (PGS78560.1),[3] LicP (WP_003186371.1),[4] NisP (WP_017864238.1),[5] ElxP (AFN69433.1),[6] CylA (WP_085405988.1),[7] AprE (WP_014417349.1),[8] CerP (AHJ59535.1),[9] PatA (AAY21150.1),[10,11] PatG (AAY21156.1),[10,11] FlaP (WP_012924050.1),[12] OphP (XPC16732.1)[13] and MpgP (WP_004993372.1),[14] and a newly identified S9 protease $WprP_2$ (WP_399537673.1) were used to generate a phylogenetic tree. A multi-sequence alignment of these proteases generated with MUSCLE was used to build a maximum-likelihood phylogenetic tree, which was visualized using the MEGA version 11.0.13.[15]

**Transformation of plasmids into *E. coli* NiCo21(DE3)**. Plasmids containing precursor and rSAM genes were obtained from Twist Bioscience and dissolved in MilliQ grade water to a final concentration of 10 ng/µL. 70 µL of *E. coli* NiCo21(DE3) electrocompetent cells were transformed in a 2 mm electroporation cuvette with either 1 µL of plasmid DNA containing the precursor gene for the precursor-only expression or 1 µL of plasmid DNA containing the precursor gene + 1 µL of plasmid DNA containing the rSAM enzyme for the coexpression of the precursor and the rSAM enzyme. The transformed cells were then grown overnight at 37 °C on LB (Miller) agar supplemented with appropriate antibiotics at a final concentration of kanamycin 30 µg/mL, spectinomycin 100 µg/mL, and chloramphenicol 25 µg/mL.

**Protein expression and purification of precursor peptides.** A colony from the transformation above was picked up by a toothpick and added to 3 mL TB medium supplemented with appropriate antibiotics in a 15 mL falcon tube. The 50 mL culture was grown overnight at 37°C and shaken at 250 rpm. The overnight culture was used to inoculate either 250 mL of antibiotic-supplemented TB media in a 1 L Erlenmeyer flask in a 1:100 (v:v) ratio. The cells were then grown at 37°C, 250 rpm until $OD_{600\ nm}$ reached 1.8-2.4. The culture was then placed on ice water for 30 min and protein expression was induced by addition of IPTG at a 0.1 mM final concentration. After induction, the culture was shaken at 16°C, 250 rpm for 18 h. The cells were collected by centrifugation at 4000 rpm for 10 min. The denaturing lysis buffer (100 mM $NaH_2PO_4$, 10 mM Tris, 8 M Urea, 10 mM imidazole, pH 8) was added to cell pellets in a ratio of 3:1 (v:w). The cell pellets were reconstituted and lysed by sonication with a Titanium 7mm solid probe (20 sec on and 15 sec off for 25 cycles at 50% amplitude). After sonication, the cell debris was removed by centrifugation at 15,000 rpm for 15 min. HisPur Ni-NTA resin (0.7 mL) was added to ~15-20 mL of supernatant in a 50 mL falcon tube and gently shaken for 1 h to allow binding of the precursor peptide to the Ni-NTA resin. Peptide-bound Ni-NTA resin was then washed with denaturing lysis buffer (2 x 1 mL for 0.7 mL resin if this buffer was used to resuspend the cell pellet), NPI-20 (50 mM $NaH_2PO_4$, 300 mM NaCl, 20 mM imidazole, pH 8, 5 x 1 mL for 0.7 mL resin) and eluted with NPI-250 (50 mM $NaH_2PO_4$, 300 mM NaCl, 250 mM imidazole, pH 8, 2.5 mL for 0.7 mL resin). Elution fractions were desalted into 50 mM Tris buffer (pH 8.0) using PD Minitrap G-10 columns. The full-length $His_6$-precursor peptide obtained by coexpression of $WprA_2B_2C_2$ was digested with GluC (10 µg per 1 mL eluate) at 37°C for 16 h and analyzed by LC-MS to detect activity from the radical SAM enzyme $WprB_2$, while the undigested full-length $His_6$-precursor peptide was used as a substrate for the biochemical characterization of S9 protease $WprP_2$.

**Protein expression and purification of S9 protease $WprP_2$.** A colony from the transformation above was picked up by a toothpick and added to 5 mL TB medium supplemented with appropriate antibiotics in a 15 mL falcon tube. The 50 mL culture was grown overnight at 37°C and shaken at 250 rpm. The overnight culture was used to inoculate either 250 mL of antibiotic-supplemented TB media in a 1 L Erlenmeyer flask in a 1:100 (v:v) ratio. The cells were then grown at 37°C, 250 rpm until $OD_{600\ nm}$ reached 1.8-2.4. The culture was then placed on ice water for 30 min and protein expression was induced by addition of IPTG at a 0.1 mM final concentration. After induction, the culture was shaken at 16°C, 250 rpm for 18 h. The cells were collected by centrifugation at 4000 rpm for 10 min. The non-denaturing buffer (50 mM $NaH_2PO_4$, 10 mM Tris, 10 mM imidazole, 300 mM NaCl, 10% glycerol, pH 8) was added to cell pellets in a ratio of 3:1 (v:w). The cell pellets were reconstituted and lysed by sonication with a Titanium 7mm solid probe (20 sec on and 15 sec off for 25 cycles at 50% amplitude). After sonication, the cell debris was removed by centrifugation at 15,000 rpm for 15 min. HisPur Ni-NTA resin (0.7 mL) was added to ~15-20 mL of supernatant in a 50 mL falcon tube and gently shaken for 1 h to allow binding of the precursor peptide to the Ni-NTA resin. Peptide-bound Ni-NTA resin was then washed with non-denaturing buffer (2 x 1 mL for 0.7 mL resin if this buffer was used to resuspend the cell pellet), NPI-20 (50 mM $NaH_2PO_4$, 300 mM NaCl, 20 mM imidazole, pH 8, 5 x 1 mL for 0.7 mL resin) and eluted with NPI-250 (50 mM $NaH_2PO_4$, 300 mM NaCl, 250 mM imidazole, pH 8, 2.5 mL for 0.7 mL resin). Elution fractions were desalted into 50 mM Tris buffer (pH 8.0) using PD Minitrap G-10 columns. Purified $His_6$-$WprP_2$ was determined by SDS-PAGE and subjected to *in vitro* assay.

**LC-MS conditions.** Data acquisition was performed using MestReNova at the following conditions.

Waters UPLC system Acquity H-class connected with Waters UV detector PDA eλ coupled with Waters TOF mass spectrometer SYNAPT G2-Si.

LC: column = Acquity BEH C18, 1.7 *μ*m, 2.1 x 50 mm; mobile phase/gradient = solvent A: $H_2O$ (+0.1% formic acid, FA), solvent B: $CH_3CN$, Initial – 95 % A, 1.5 min – 40 % A, 2.5 min – 2 % A, 3.5 min – 2 % A, 3.7 min – 95 % A, 6 min – 95 % A; flow rate = 0.4 mL/min; column temp. = 40 °C; Injection volume = 1 μL, Run time = 6 min

MS: polarity = positive; capillary voltage = +0.7 kV; cone voltage = 40.0 V; source temperature = 120 °C; desolvation temperature = 500 °C; cone gas flow = 30 L/h; desolvation gas flow = 800 L/h; collision energy = 4 V; mass range = *m/z* 250–2000; scan duration = 0.5 s; interscan delay = 0.025 s; data acquisition = continuum mode; Lockspray (Leucine enkephalin); scan duration = 1.0 s; interval = 10 scans.

MS/MS: polarity = positive; scan duration = 1.0 s; inter-scan delay = 0.025 s; data acquisition = continuum mode; mass range and collision energy are specified in the respective figure legends.

***In vitro* assay of S9 protease $WprP_2$.** $His_6$-$WprP_2$ biochemical characterization was carried out in 50 mM Tris-HCl buffer (pH 8.0) with purified full-length $His_6$-precursor peptide (0.1 - 0.2 mM), 3 mM of dithiothreitol (DTT) and 5 uM of enzyme $His_6$-$WprP_2$ in a reaction volume of 100 uL. For the negative control, enzyme $His_6$-$WprP_2$ was boiled by heating at 80 °C for 10 min. The reaction solution was incubated at 25 °C for 2 h and quenched with absolute methanol. After centrifugation at 15,000 rpm for 10 min, the products were then subjected to LC-MS analysis as mentioned above.

**Mutation on $WprA_1$, $WprA_2$ and $WprP_2$.** The mutation at R33G and N36Q were introduced to precursor peptide $WprA_1$ with primers listed in Table S4. The mutation at Q44A, H38A and R59A were introduced to precursor peptide $WprA_2$ with primers listed in Table S4. The mutation at H621A, D590A and S507A were introduced to S9 protease $WprP_2$ with primers listed in Table S4. These plasmids containing mutated residue were amplified with Phusion high-fidelity DNA polymerase (New England Biolabs) in PCR thermal cycler (Eppendorf), followed by digestion with enzyme DpnI (New England Biolabs), gel purification with DNA extraction kit (Bio Basic), DNA 5' phosphorylation with T4 polynucleotide kinase (New England Biolabs), and DNA ligation with T4 DNA ligase (New England Biolabs). The cloned plasmids were transformed into *E. coli* DH5$\alpha$, then harvested with Monarch plasmid miniprep kit (New England Biolabs), and sequence verified at Genewiz from Azenta Life Sciences.

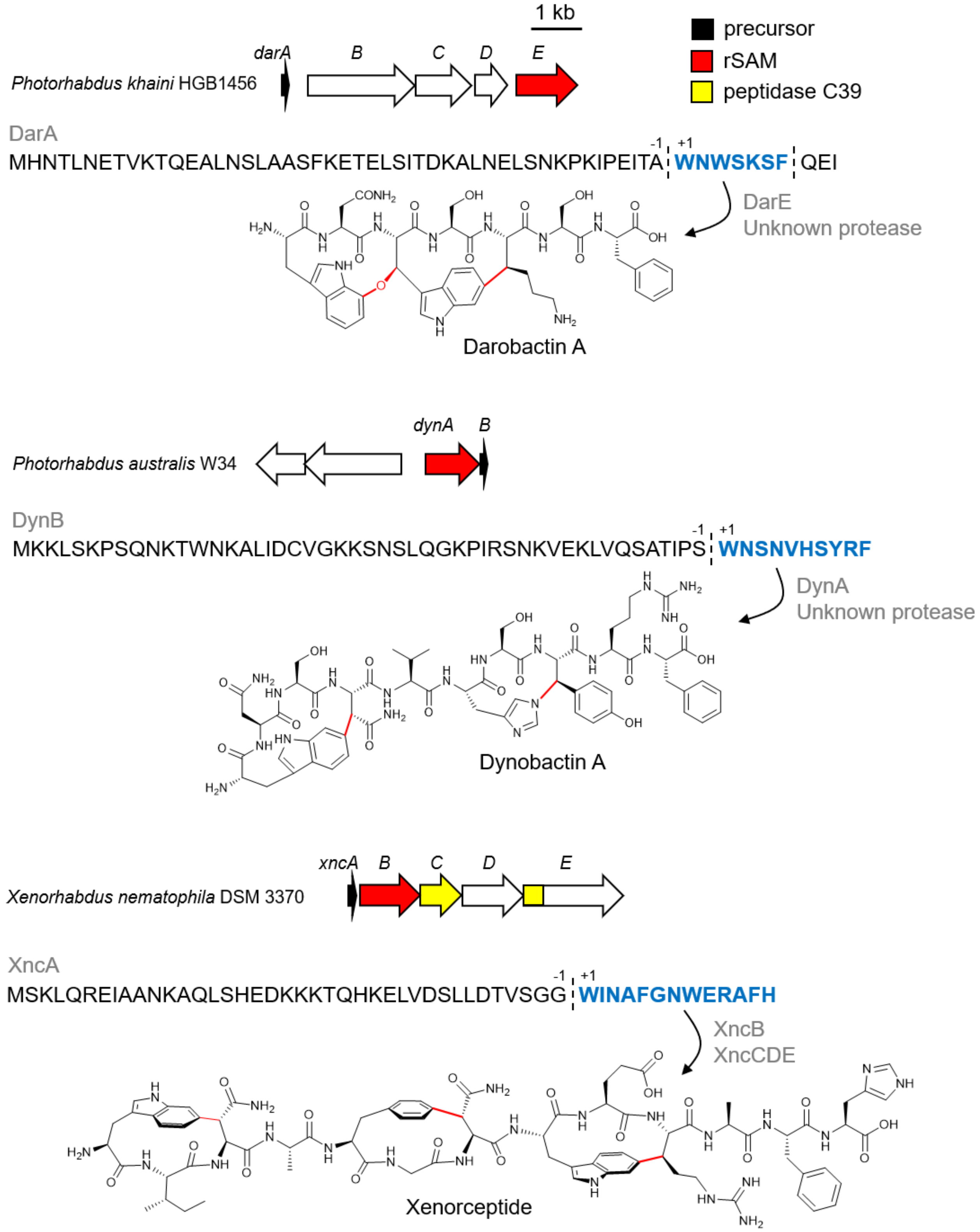


**Figure S1.** Proteases involved in the rSAM-RiPP biosynthesis of darobactin A,[16,17] dynobactin A,[18] and xenorceptide.[19] Protease cleavage sites are shown as dashed lines. Core peptides are shown as blue colored bold letters. Residues within the precursor sequence are numbered '+1' from the start of the core peptide.

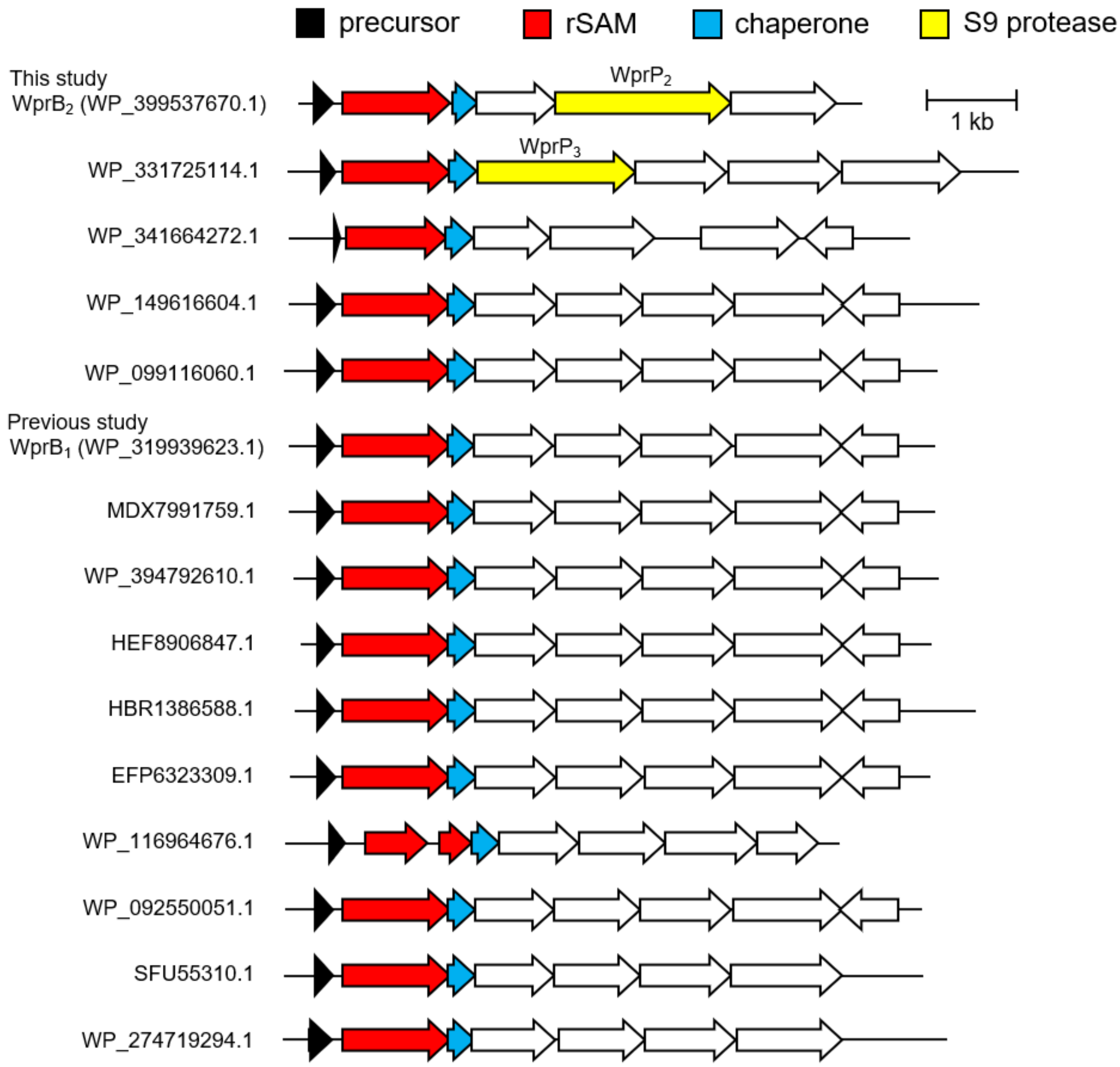


**Figure S2.** BGCs of 16 homologous WprB whose putative precursor peptides contain one or three repeated WPR motifs. Two BGCs encoded S9 proteases, designated as $WprP_2$ (WP_399537673.1) and $WprP_3$ (WP_331725116.1).

WprA$_2$: MTDFQPSFETADTDVLR**WPR**HTGEDFQPAFEATDTDTLR**WPR**HTGDDFQPAFEATDTDTLR**WPR**HTE

His$_6$-WprA$_2$ —Ni-affinity / GluC→

**1a** TADTDVLR**WPR**HTGE

**2a** ATDTDTLR**WPR**HTGDDFQPAFE

**3a** ATDTDTLR**WPR**HTE

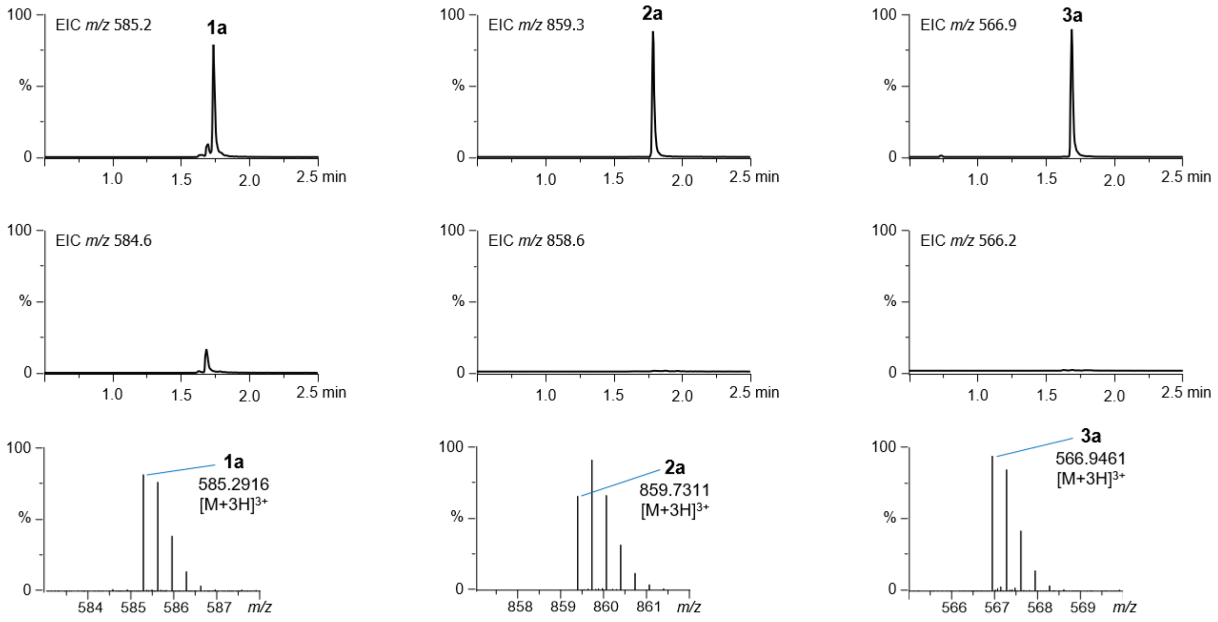


**Figure S3.** *In vivo* coexpression of His$_6$-WprA$_2$ followed by Ni-affinity purification and GluC digestion yielded fragments **1a**-**3a**. The EIC chromatogram and MS spectra of fragments **1a**-**3a**.

$WprA_2$: MTDFQPSFETADTDVLR**WPR**HTGEDFQPAFEATDTDTLR**WPR**HTGDDFQPAFEATDTDTLR**WPR**HTE

$His_6$-$WprA_2$ + $WprB_2$ —(Ni-affinity / GluC)→

**1a** TADTDVLR**WPR**HTGE

**2a** ATDTDTLR**WPR**HTGDDFQPAFE

**3a** ATDTDTLR**WPR**HTE

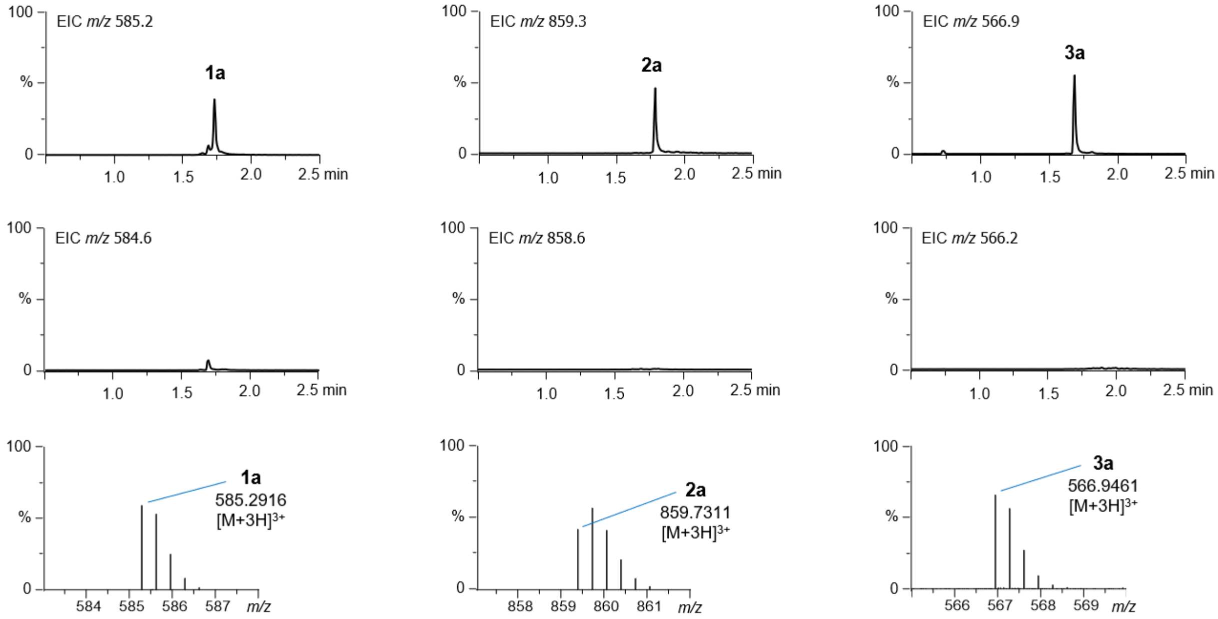


**Figure S4.** *In vivo* coexpression of $His_6$-$WprA_2$ + $WprB_2$ followed by Ni-affinity purification and GluC digestion yielded fragments **1a**-**3a**. The EIC chromatogram and MS spectra of fragments **1a**-**3a**.

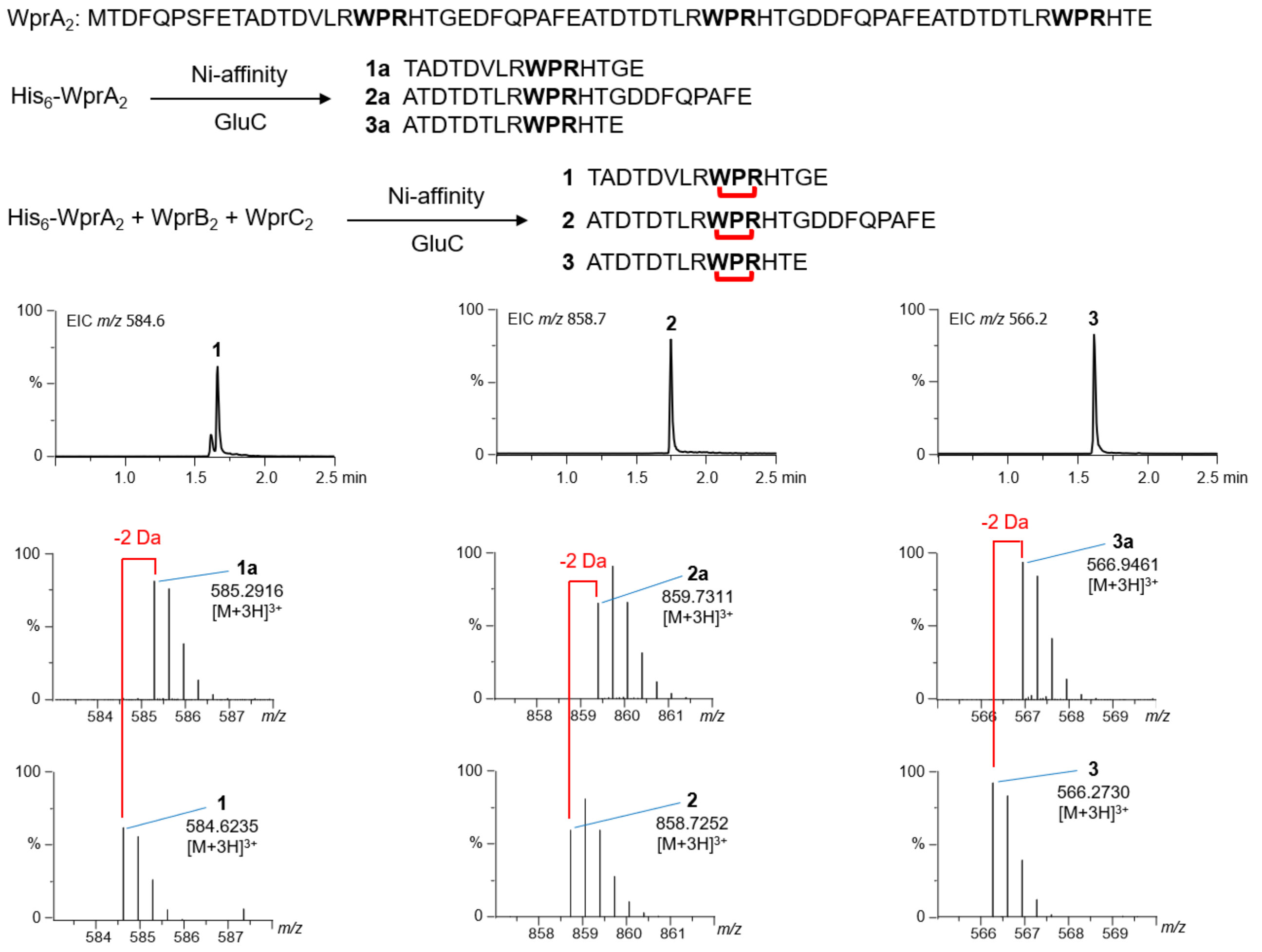


**Figure S5.** *In vivo* coexpression of His$_6$-WprA$_2$ + WprB$_2$ + WprC$_2$ followed by Ni-affinity purification and GluC digestion yielded fragments **1**-**3**. The EIC chromatogram and MS spectra of fragments **1**-**3**. Cross-link formation on the peptide sequences is shown as red connectors.

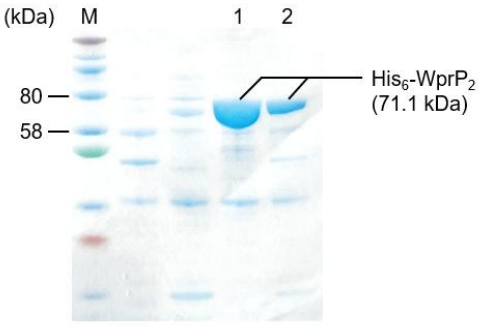


**Figure S6.** SDS-PAGE of WprP$_2$. M: molecular weight marker. 1: *in vivo* expression of His$_6$-WprP$_2$ in *E. coli* NiCo21 (DE3) and via Ni-affinity purification, 10 µL loaded into the well. 2: *in vivo* expression of His$_6$-WprP$_2$ in *E. coli* NiCo21 (DE3) and via Ni-affinity purification, 5 µL loaded into the well. Theoretical molecular weight of recombinant His$_6$-WprP$_2$ is 71.1 kDa.

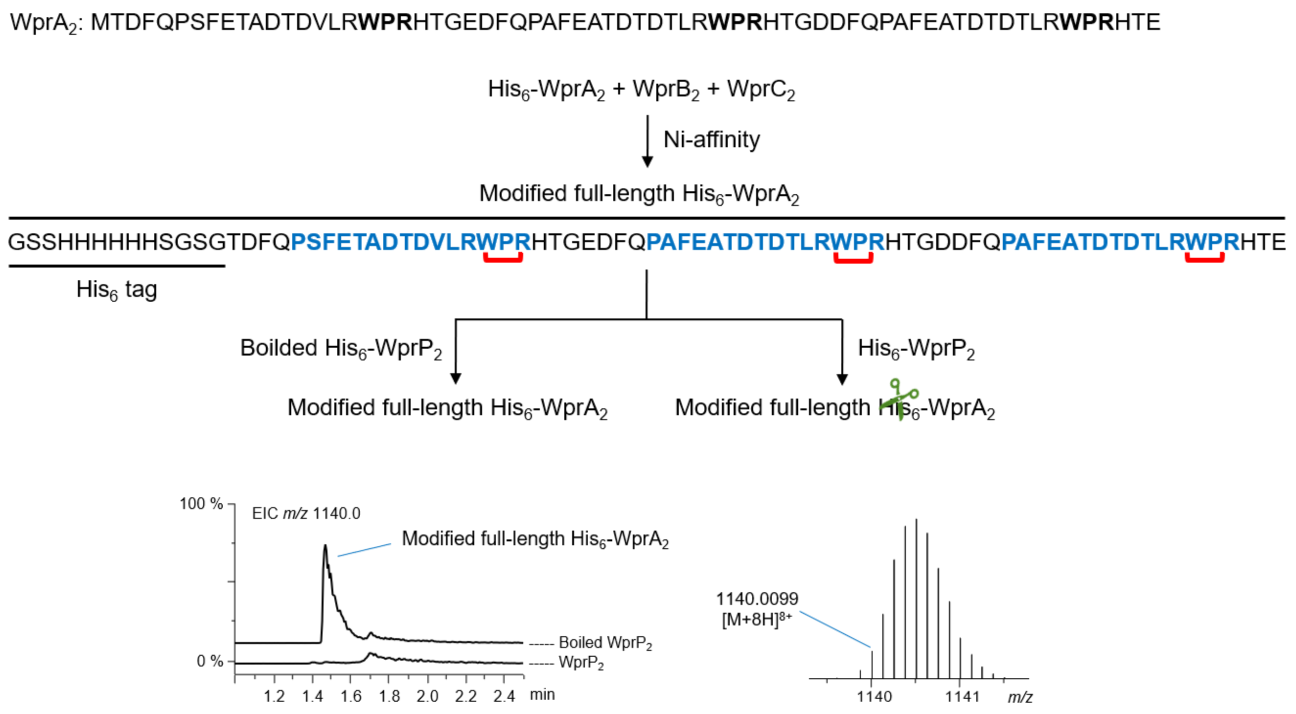


**Figure S7.** *In vitro* assay of His$_6$-WprP$_2$ or boiled His$_6$-WprP$_2$ + modified full-length His$_6$-WprA$_2$. The EIC chromatogram and MS spectra of modified full-length His$_6$-WprA$_2$. The presence of modified full-length His$_6$-WprA$_2$ was only found in the reaction with boiled enzyme. Cross-link formation on the peptide sequences is shown as red connectors. Core peptides are shown as blue colored bold letters.

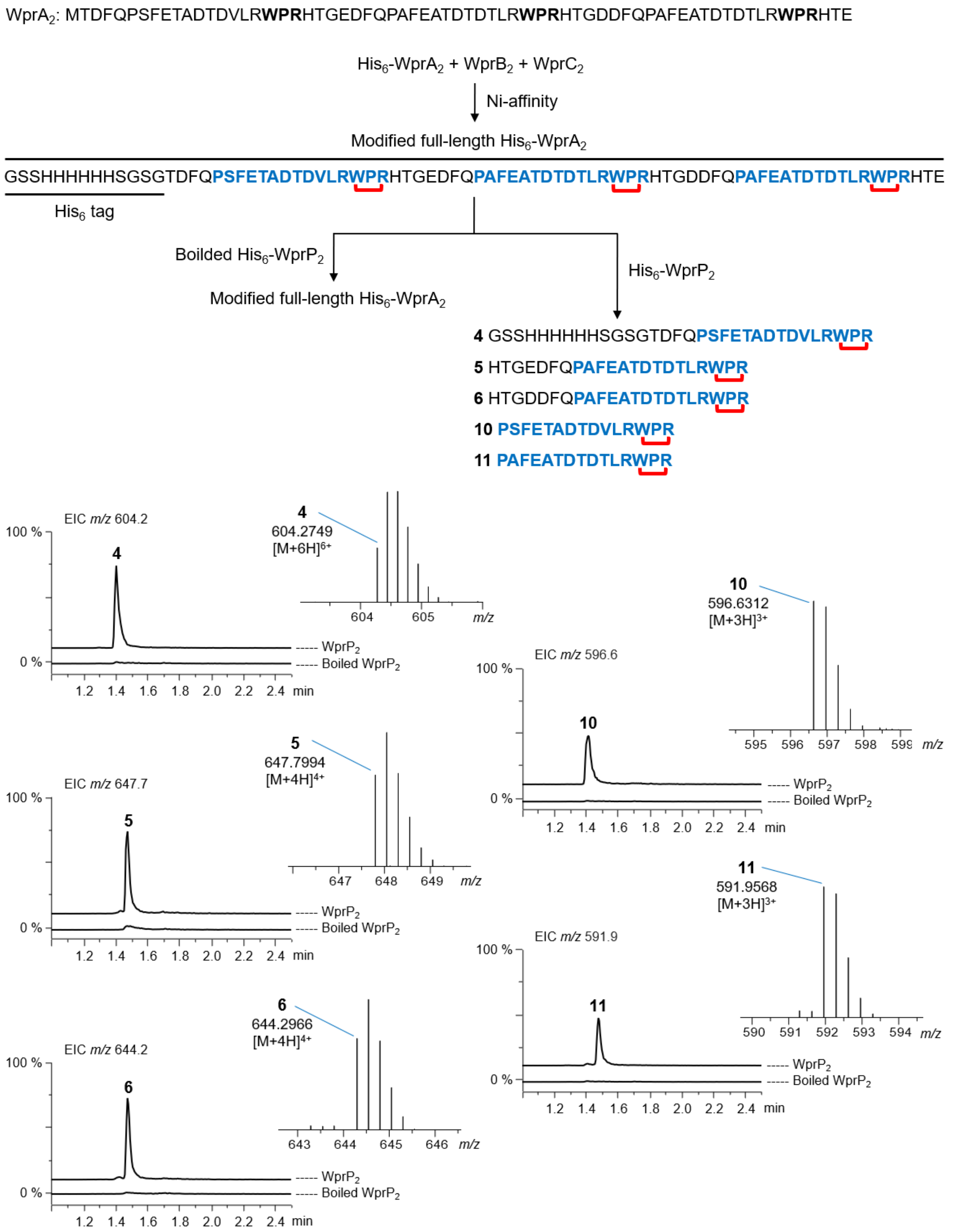


**Figure S8.** *In vitro* assay of His$_6$-WprP$_2$ or boiled His$_6$-WprP$_2$ + modified full-length His$_6$-WprA$_2$. The EIC chromatogram and MS spectra of fragments **4-6** and **10-11**. The first cleavage after WPR motif catalyzed by WprP$_2$ generates fragments **4-6**, and second cleavage before the Pro amino acid at the 12th residue preceding the WPR motif generates fragments **10-11**. Cross-link formation on the peptide sequences is shown as red connectors. Core peptides are shown as blue colored bold letters.

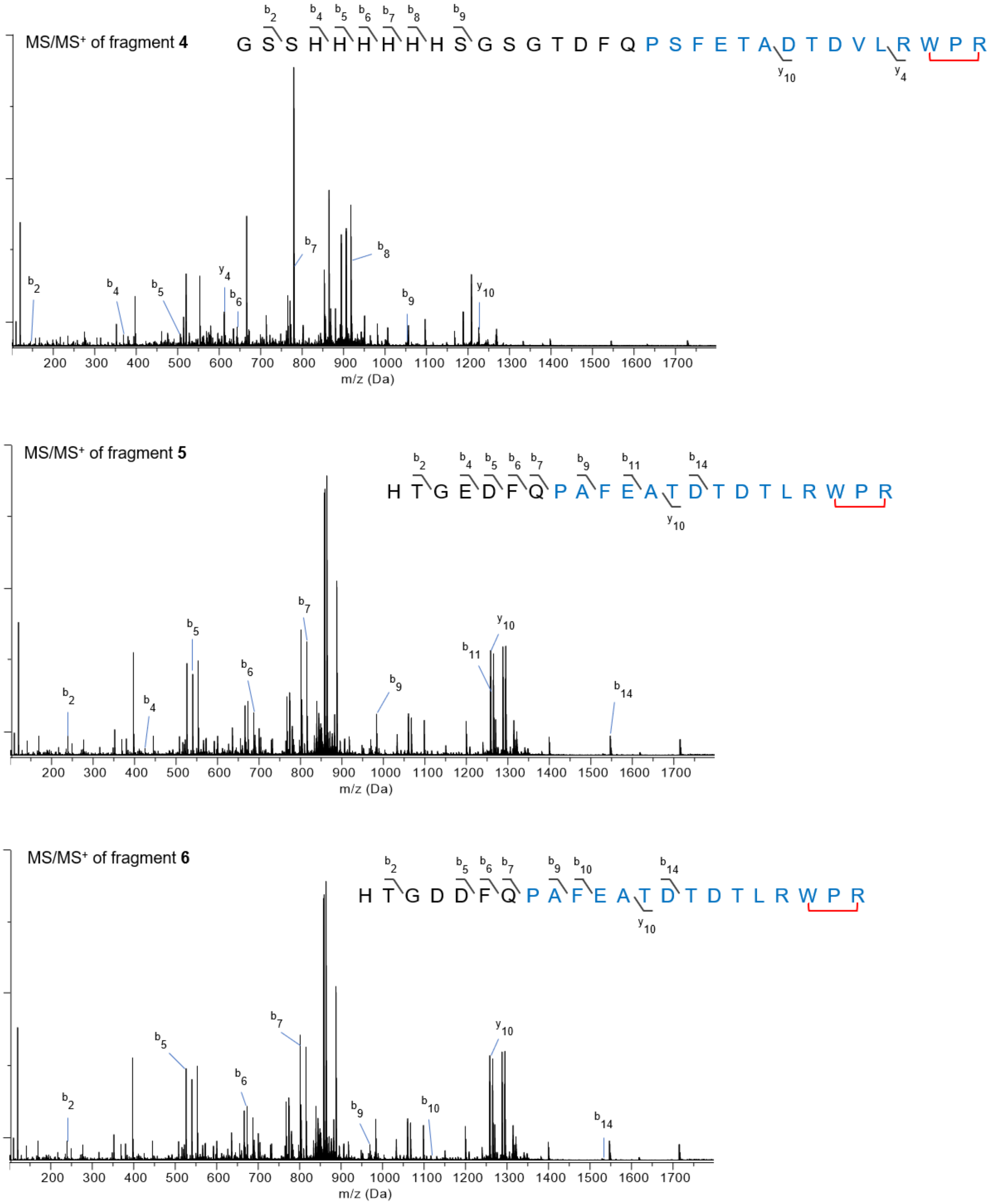


**Figure S9.** The MS/MS spectra of fragments **4-6**.

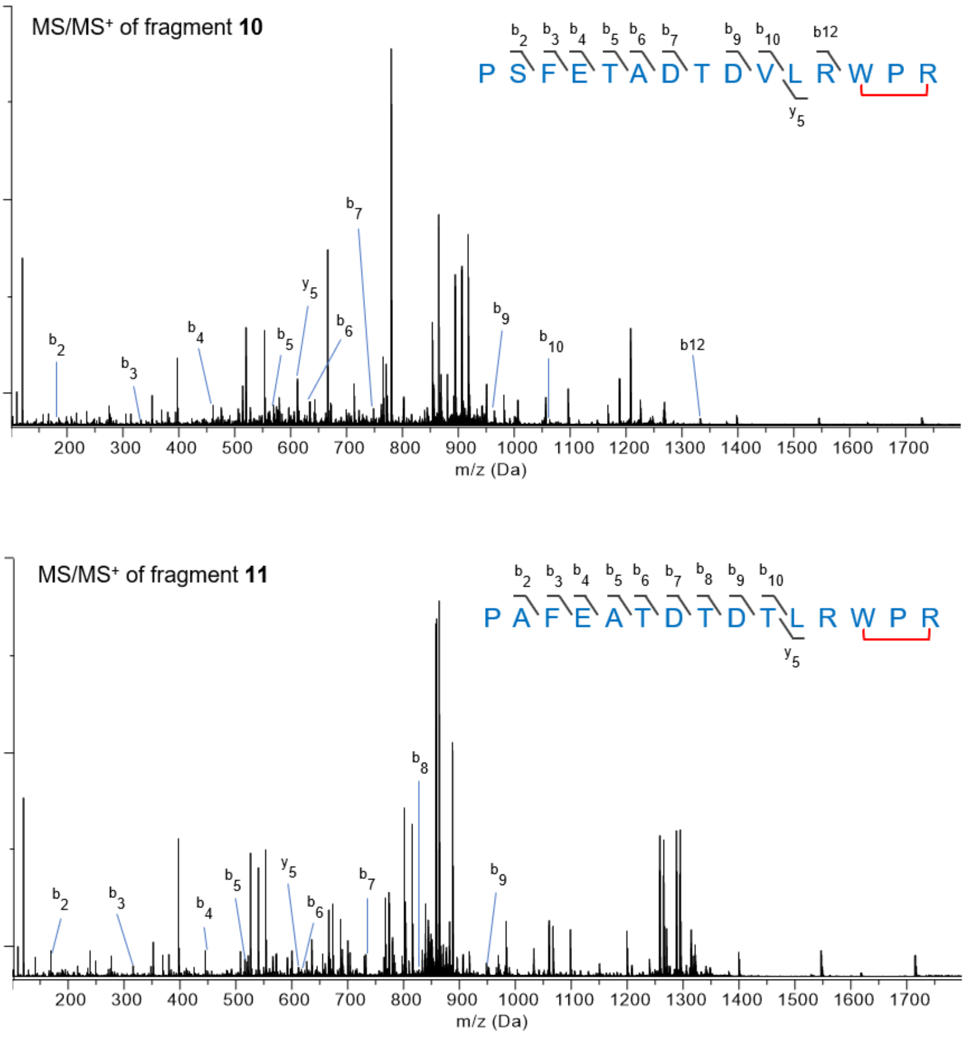


**Figure S10.** The MS/MS spectra of fragments **10** and **11**.

$WprA_2$: MTDFQPSFETADTDVLR**WPR**HTGEDFQPAFEATDTDTLR**WPR**HTGDDFQPAFEATDTDTLR**WPR**HTE

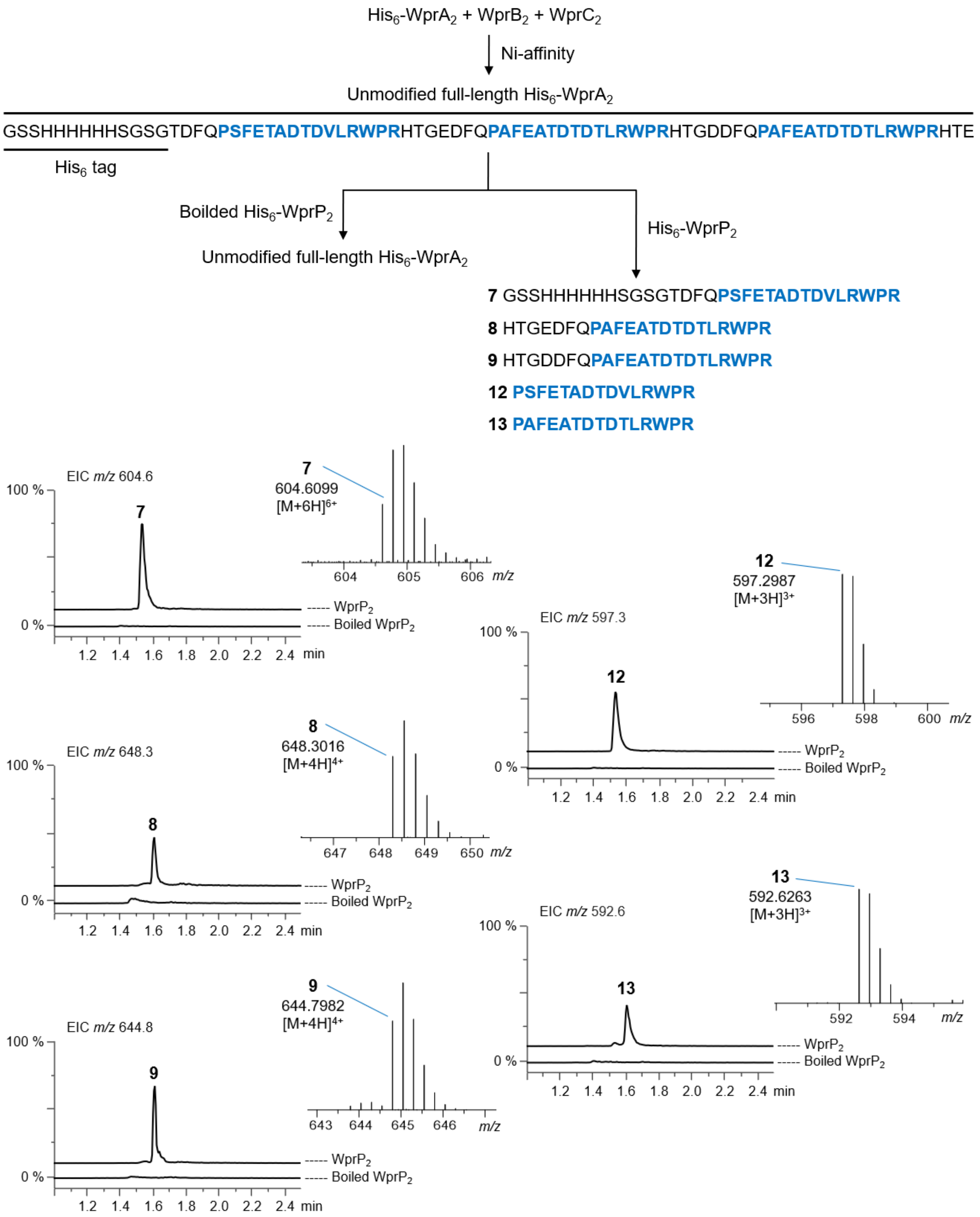


**Figure S11.** *In vitro* assay of $His_6$-$WprP_2$ or boiled $His_6$-$WprP_2$ + unmodified full-length $His_6$-$WprA_2$. The EIC chromatogram and MS spectra of fragments **7-9** and **12-13**. The first cleavage after WPR motif catalyzed by $WprP_2$ generates fragments **7-9**, and second cleavage before the Pro amino acid at the 12$^{th}$ residue preceding the WPR motif generates fragments **12-13**. Cross-link formation on the peptide sequences is shown as red connectors. Core peptides are shown as blue colored bold letters.

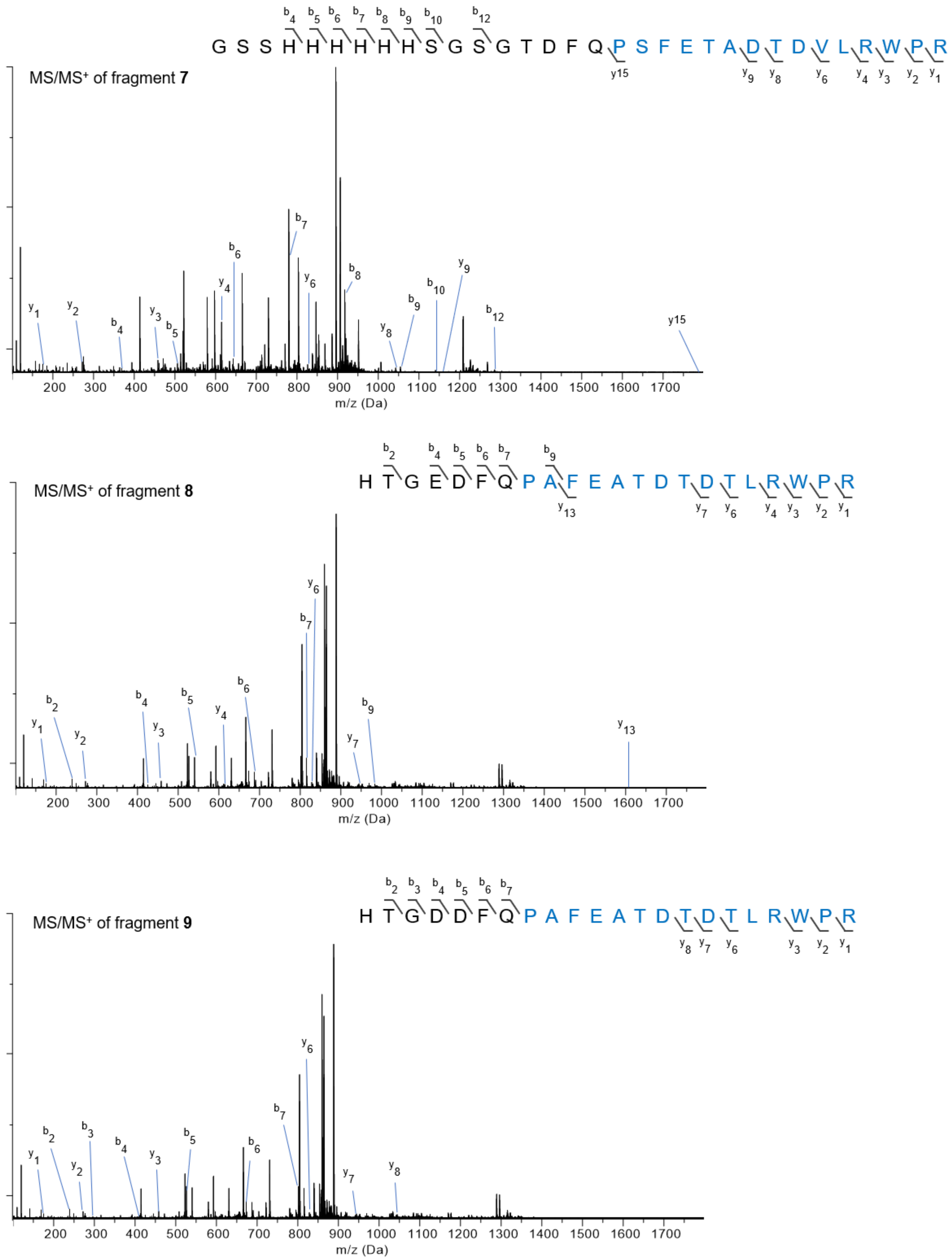


**Figure S12.** The MS/MS spectra of fragments **7-9**.

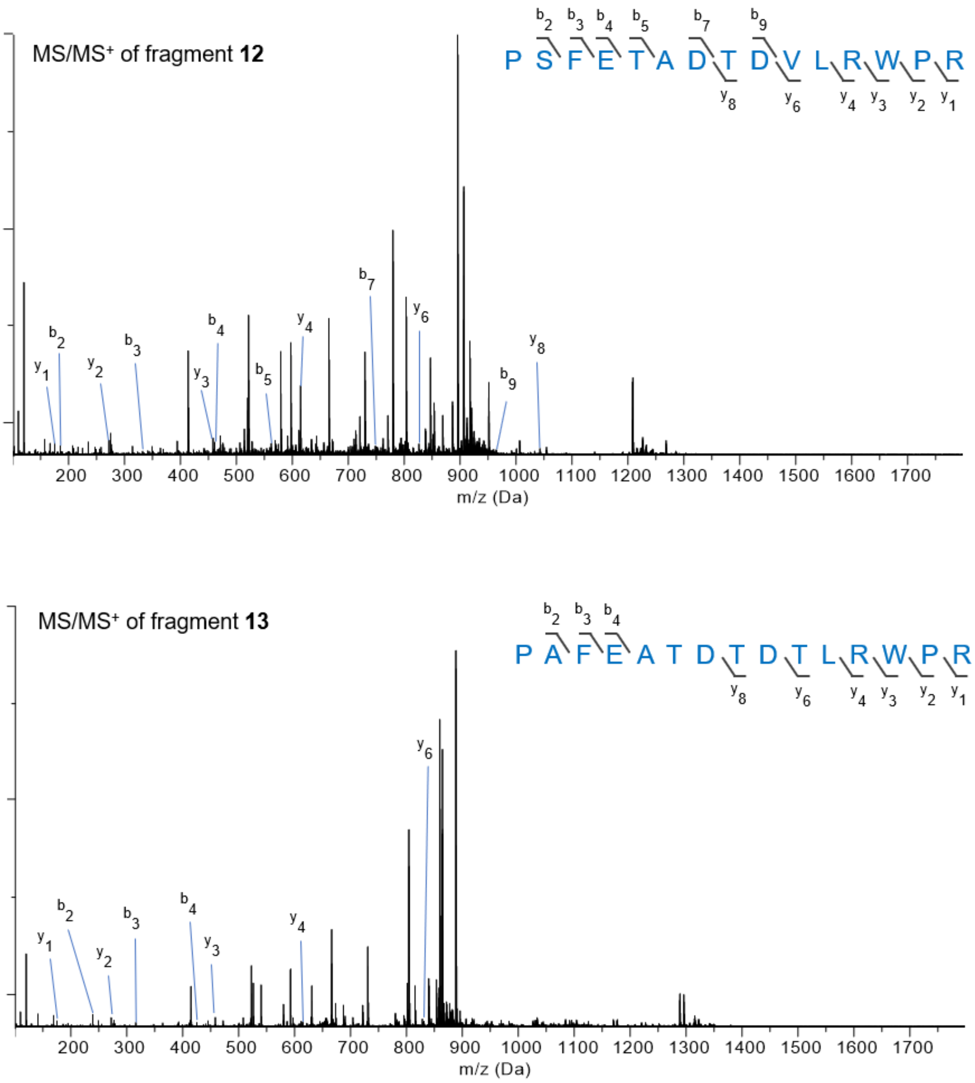


**Figure S13.** The MS/MS spectra of fragments **12** and **13**.

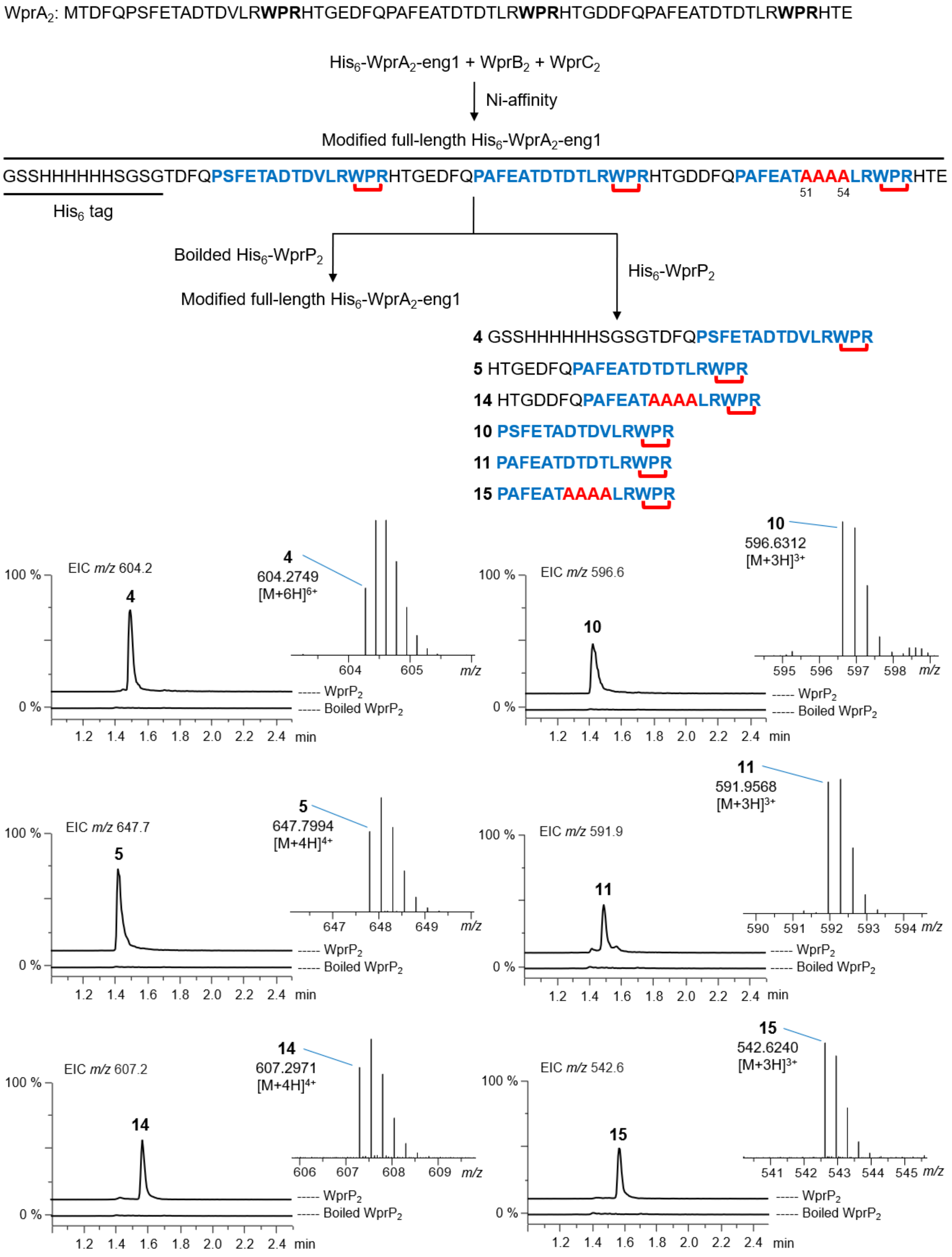


**Figure S14.** *In vitro* assay of His$_6$-WprP$_2$ or boiled His$_6$-WprP$_2$ + modified full-length His$_6$-WprA$_2$-eng1. The EIC chromatogram and MS spectra of fragments **4**, **5**, **10**, **11**, **14** and **15**. Multiple mutation at D51A/T52A/D53A/T54A on the precursor peptide did not affect cleavage activity. Core peptides and mutated residues are shown as blue and red colored bold letters, respectively.

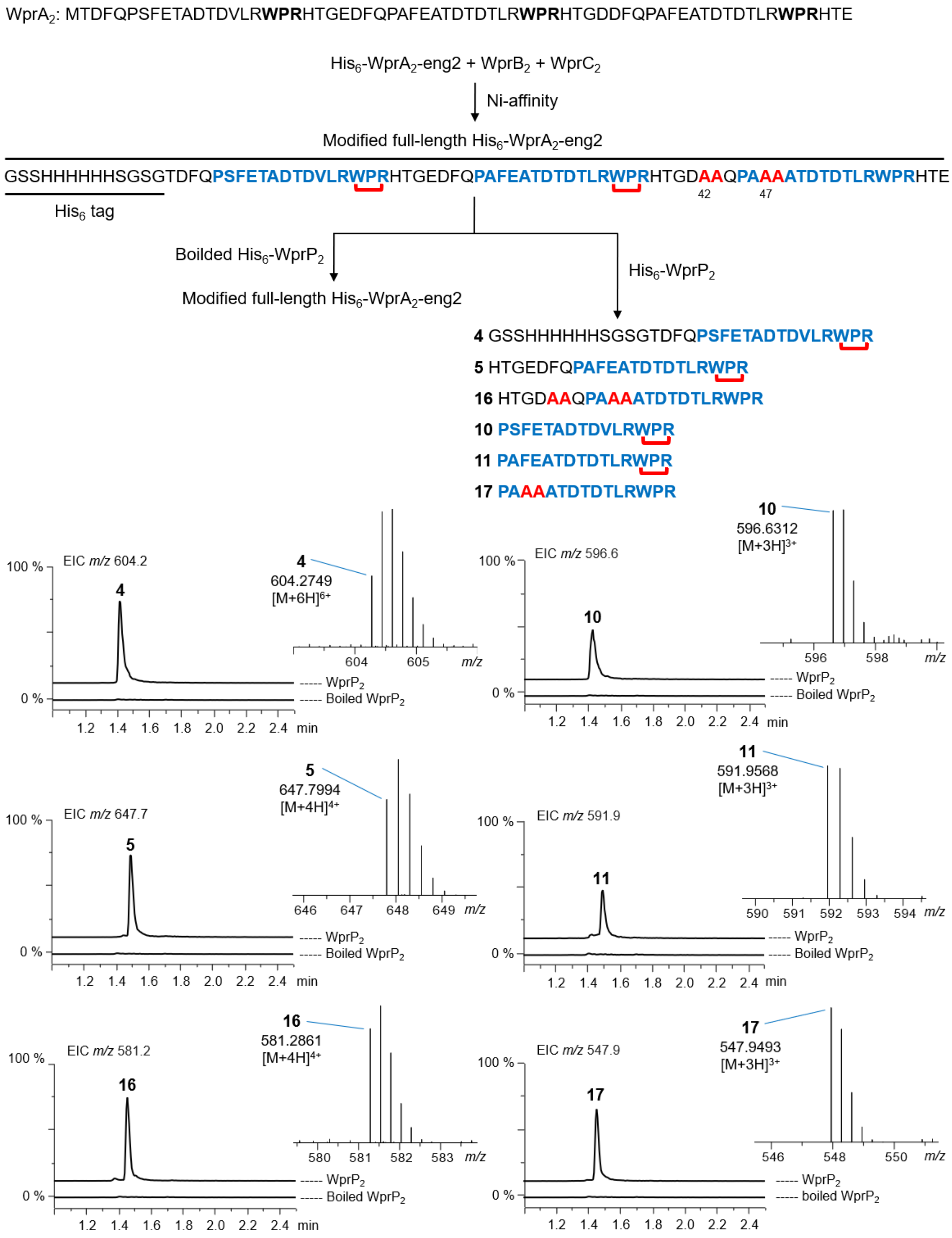


**Figure S15.** *In vitro* assay of His$_6$-WprP$_2$ or boiled His$_6$-WprP$_2$ + modified full-length His$_6$-WprA$_2$-eng2. The EIC chromatogram and MS spectra of fragments **4**, **5**, **10**, **11**, **16** and **17**. Multiple mutation at D42A/F43A/P47A/E48A on the precursor peptide did not affect cleavage activity. Core peptides and mutated residues are shown as blue and red colored bold letters, respectively.

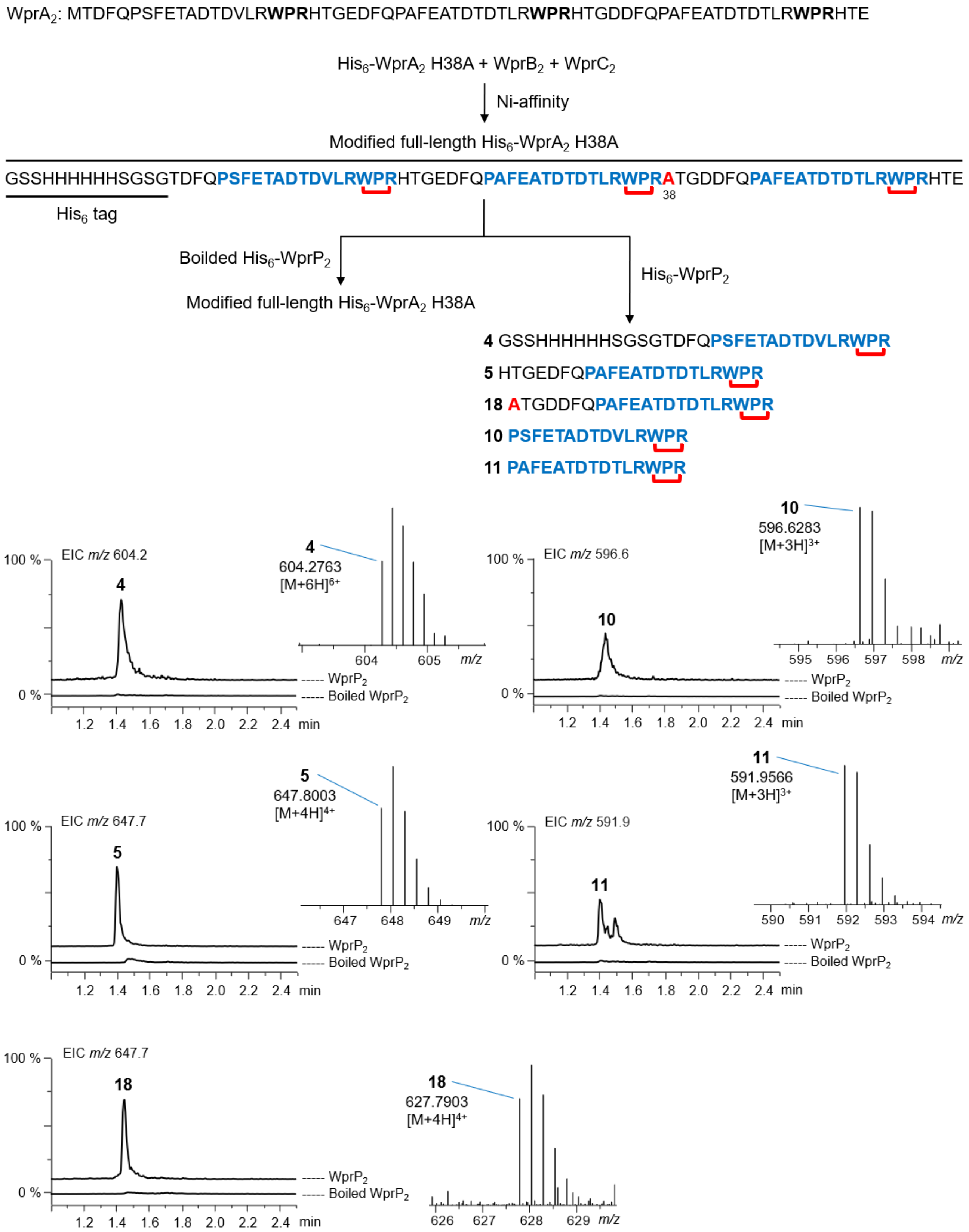


**Figure S16.** *In vitro* assay of $His_6$-$WprP_2$ or boiled $His_6$-$WprP_2$ + modified full-length $His_6$-$WprA_2$ H38A. The EIC chromatogram and MS spectra of fragments **4**, **5**, **10**, **11** and **18**. A single mutation at H38A on the precursor peptide did not affect cleavage activity. Core peptides and mutated residues are shown as blue and red colored bold letters, respectively.

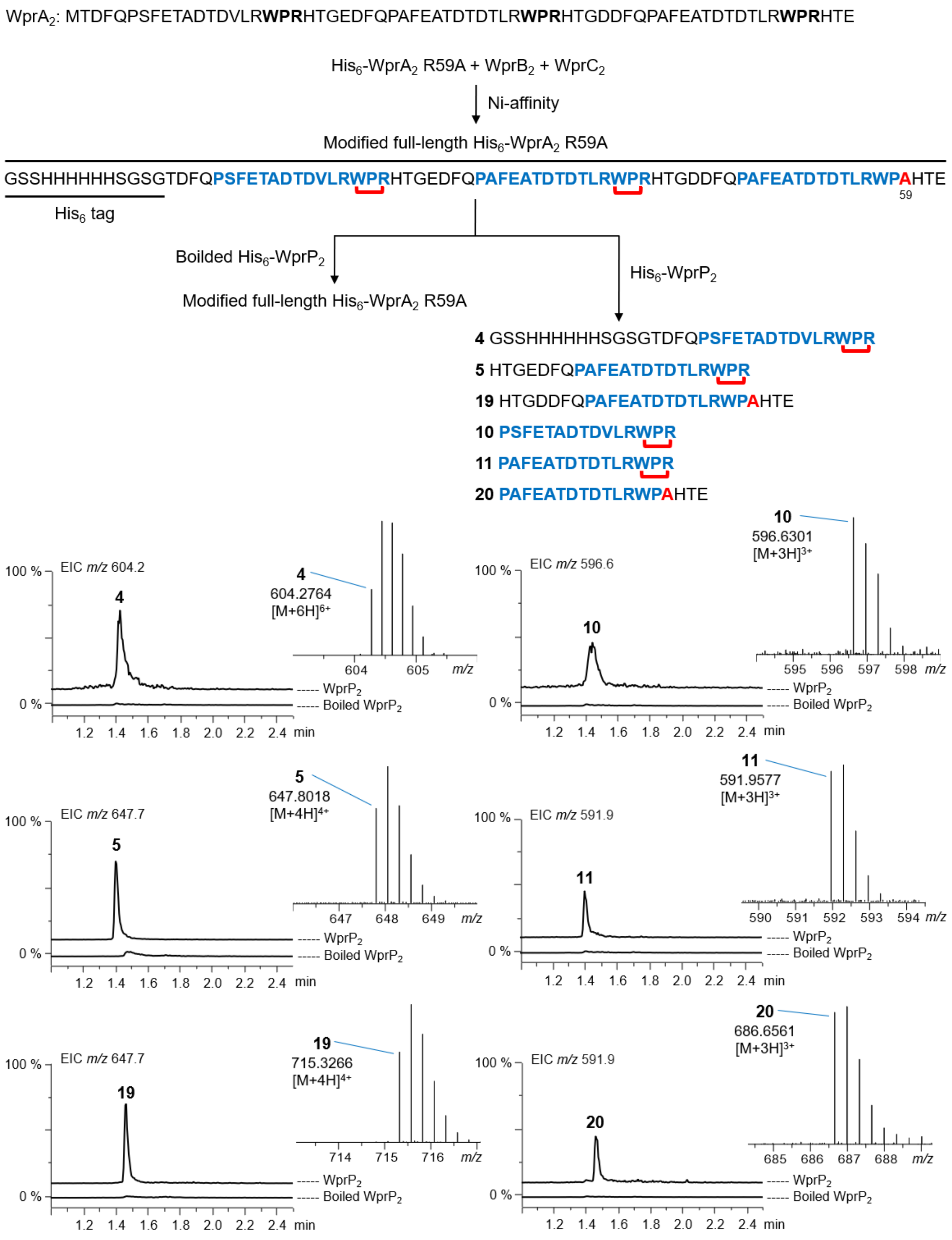


**Figure S17.** *In vitro* assay of His$_6$-WprP$_2$ or boiled His$_6$-WprP$_2$ + modified full-length His$_6$-WprA$_2$ R59A. The EIC chromatogram and MS spectra of fragments **4**, **5**, **10**, **11**, **19** and **20**. A single mutation at R59A on the precursor peptide prevented cleavage at this position. Core peptides and mutated residues are shown as blue and red colored bold letters, respectively.

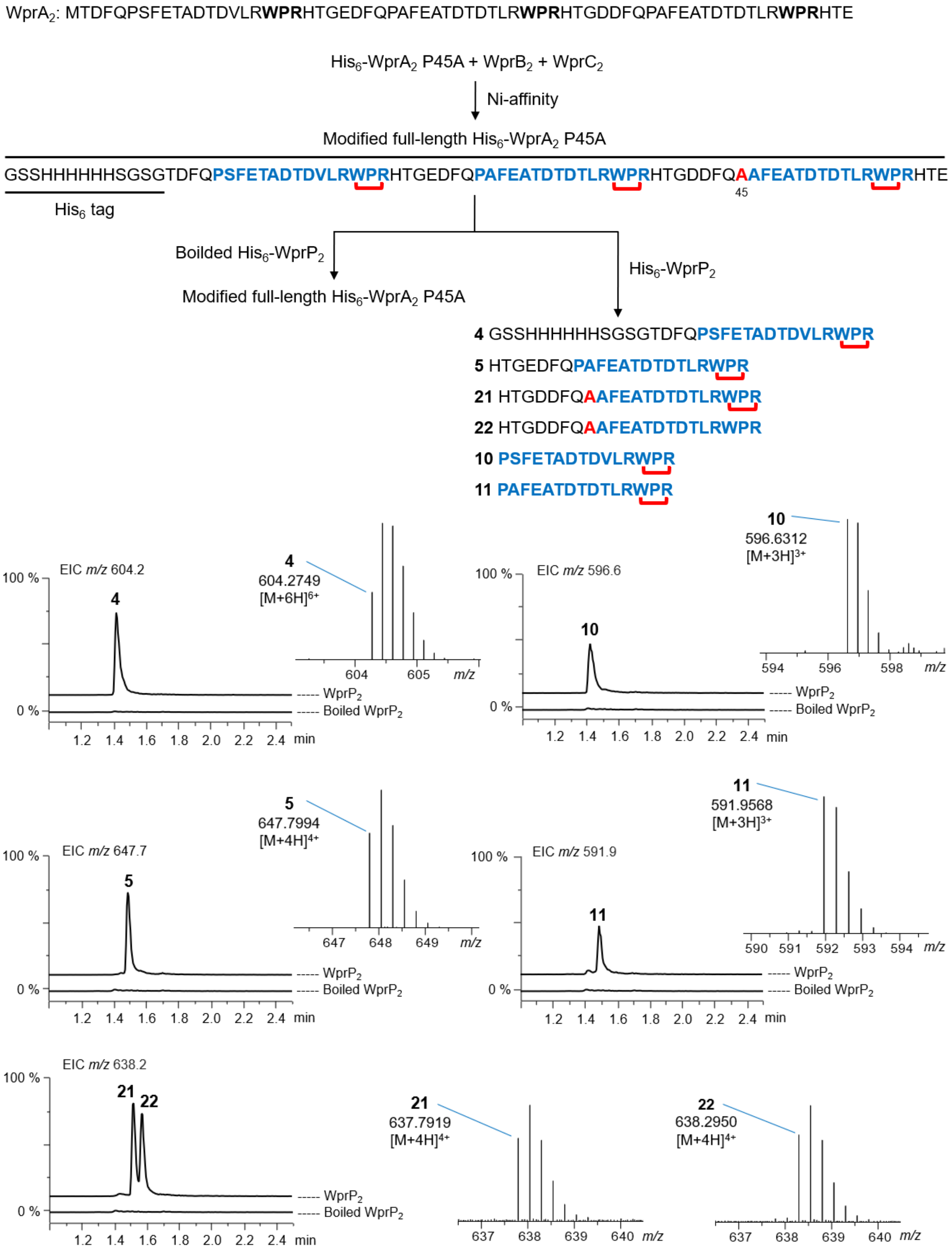


**Figure S18.** *In vitro* assay of His$_6$-WprP$_2$ or boiled His$_6$-WprP$_2$ + modified full-length His$_6$-WprA$_2$ P45A. The EIC chromatogram and MS spectra of fragments **4**, **5**, **10**, **11**, **21** and **22**. A single mutation at P45A on the precursor peptide prevented cleavage at this position. Core peptides and mutated residues are shown as blue and red colored bold letters, respectively.

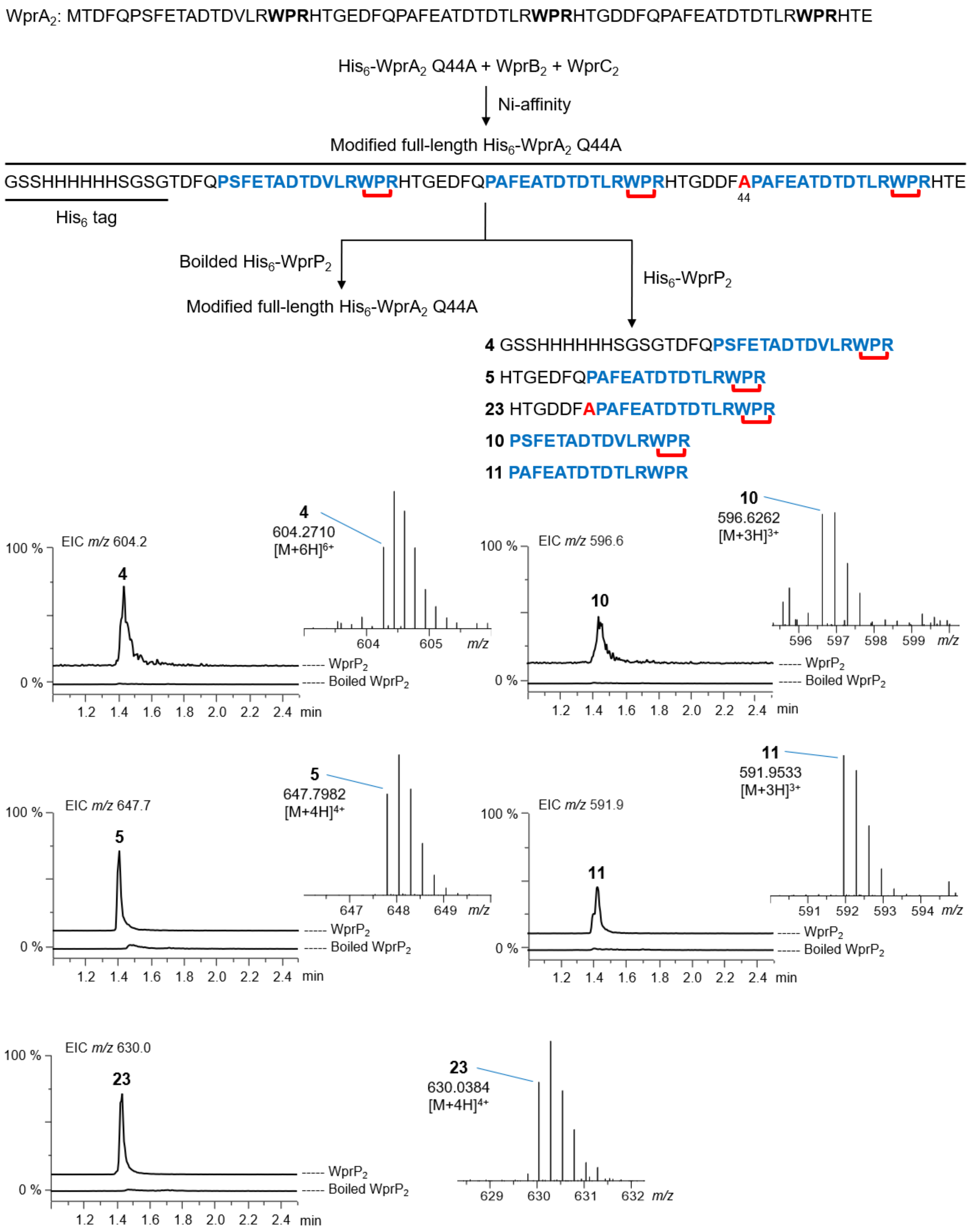


**Figure S19.** *In vitro* assay of His$_6$-WprP$_2$ or boiled His$_6$-WprP$_2$ + modified full-length His$_6$-WprA$_2$ Q44A. The EIC chromatogram and MS spectra of fragments **4**, **5**, **10**, **11** and **23**. A single mutation at Q44A on the precursor peptide prevented cleavage at this position. Core peptides and mutated residues are shown as blue and red colored bold letters, respectively.

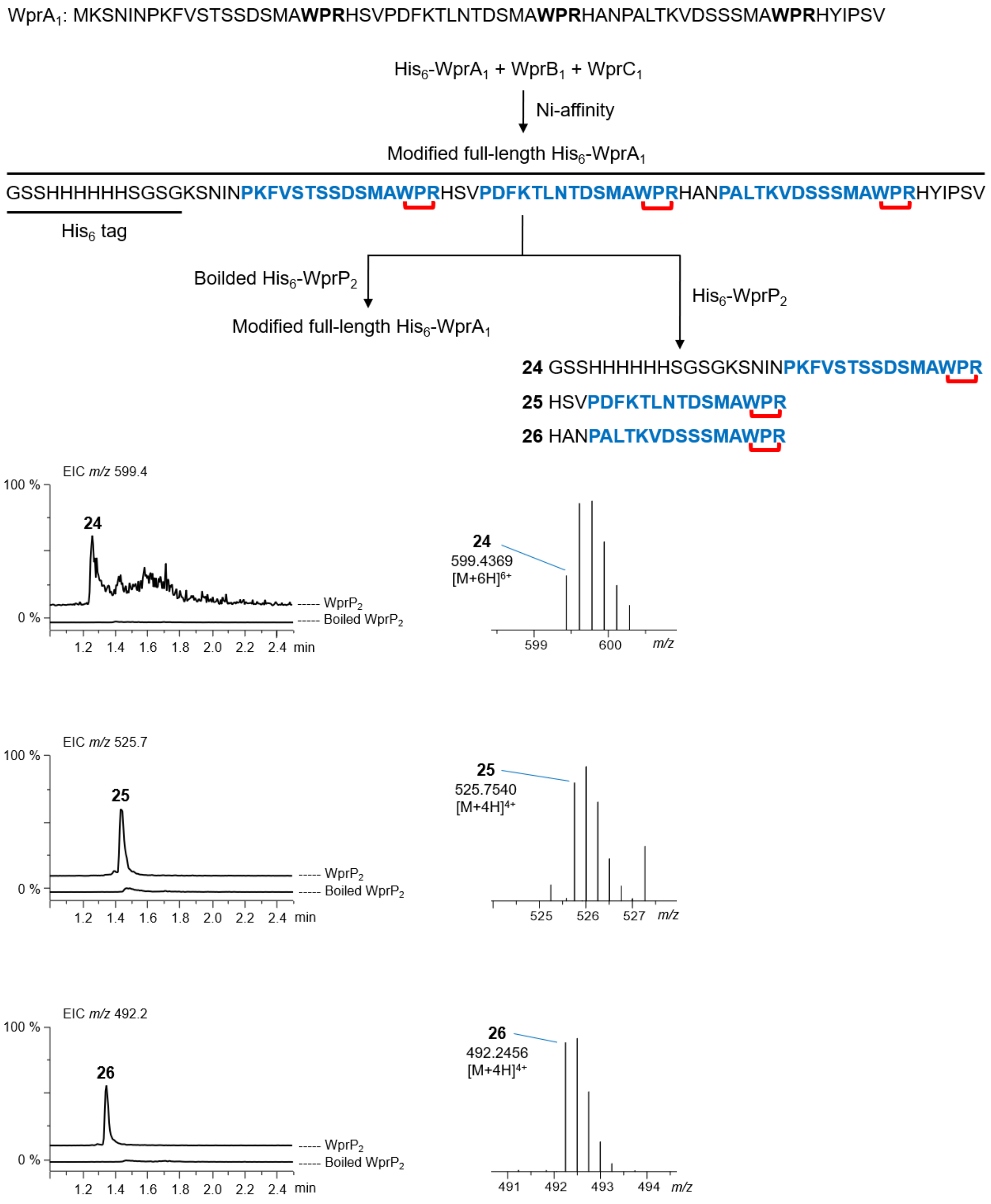


**Figure S20.** *In vitro* assay of His$_6$-WprP$_2$ or boiled His$_6$-WprP$_2$ + modified full-length His$_6$-WprA$_1$. The EIC chromatogram and MS spectra of fragments **24-26**. The first cleavage after WPR motif catalyzed by WprP$_2$ generates fragments **24-26**, however second cleavage was not detected. Cross-link formation on the peptide sequences is shown as red connectors. Core peptides are shown as blue colored bold letters.

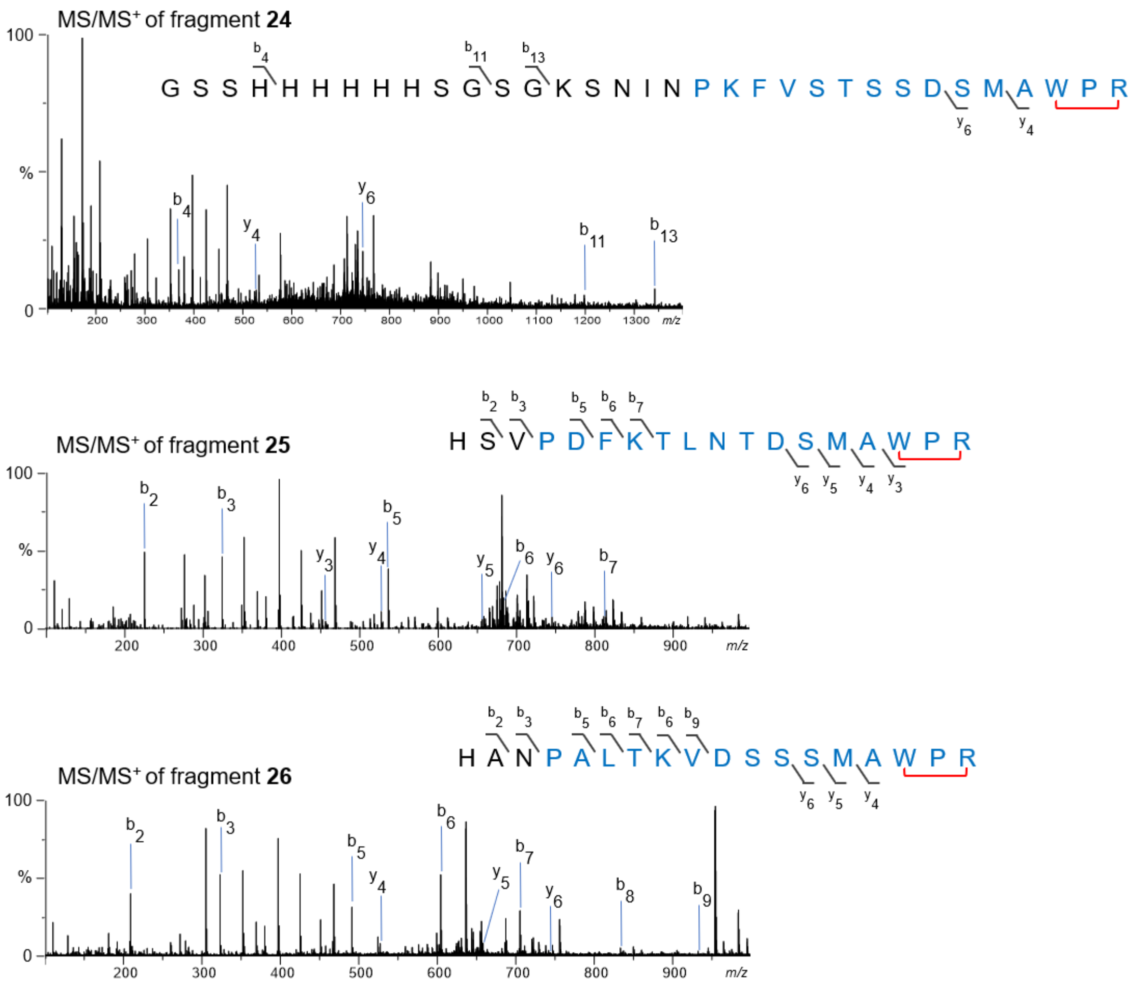


**Figure S21.** The MS/MS spectra of fragments **24-26**.

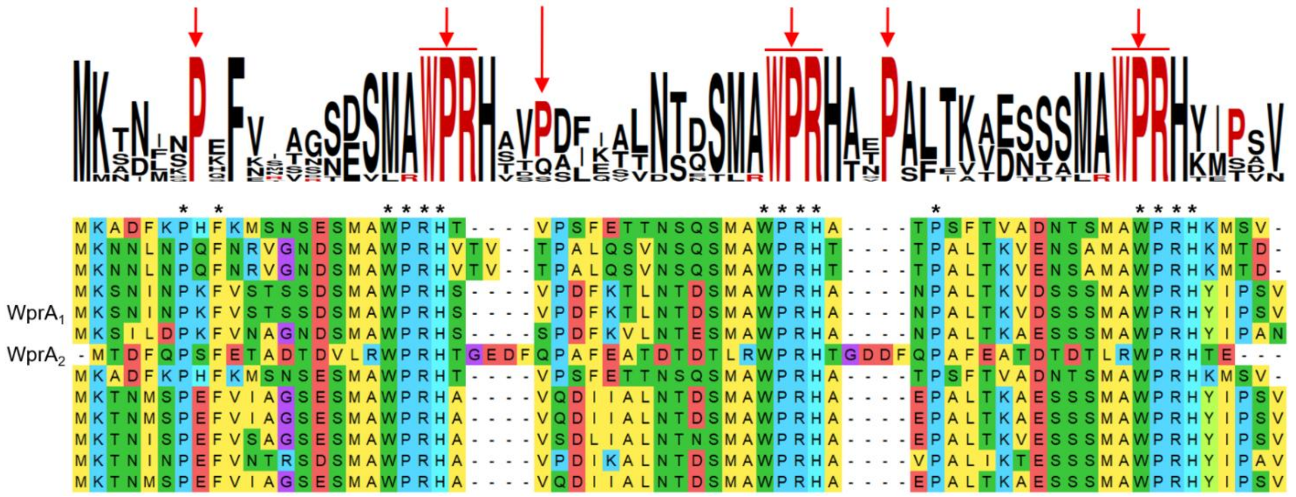


**Figure S22.** A sequence alignment generated using CLUSTAL W.[20] Comparison of precursor peptides WprA that contain three repeated WPR motifs. Conserved residues on the precursor sequences are shown as asterisk symbols.

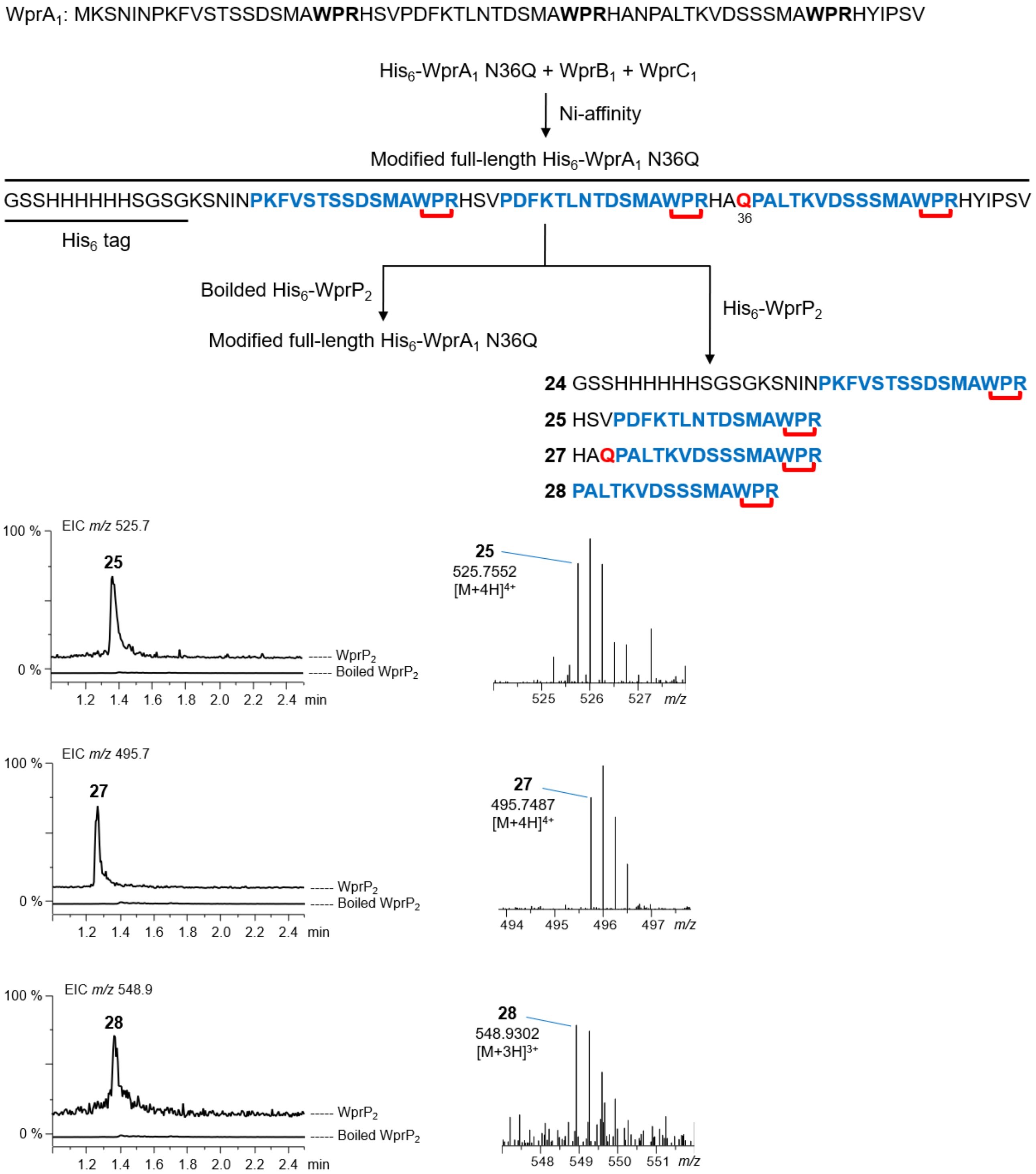


**Figure S23.** *In vitro* assay of His$_6$-WprP$_2$ or boiled His$_6$-WprP$_2$ + modified full-length His$_6$-WprA$_1$ N36Q. The EIC chromatogram and MS spectra of fragments **25**, **27** and **28**. WprP$_2$ catalyzes the first cleavage after WPR motif to generate fragments **25** and **27**, and a single mutation at N36Q on the precursor peptide allows a second cleavage before the Pro amino acid at the 12$^{th}$ residue preceding the WPR motif generates fragment **28**. Core peptides and mutated residues are shown as blue and red colored bold letters, respectively.

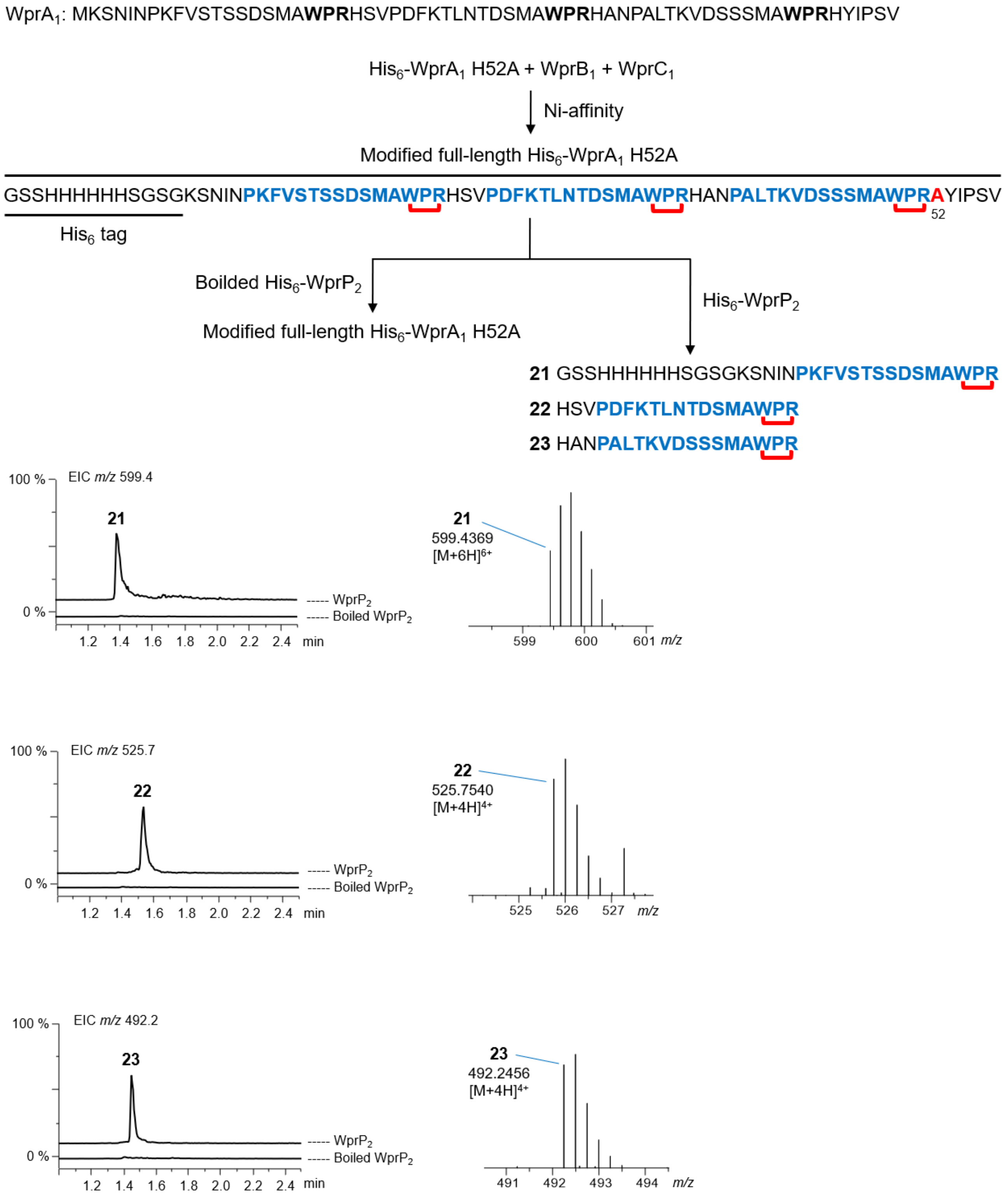


**Figure S24.** *In vitro* assay of $His_6$-$WprP_2$ or boiled $His_6$-$WprP_2$ + modified full-length $His_6$-$WprA_1$ H52A. The EIC chromatogram and MS spectra of fragments **21**-**23**. $WprP_2$ catalyzes the first cleavage after WPR motif to generate fragments **21**-**23**. A single mutation at H52A on the precursor peptide did not prevent cleavage at this position. Core peptides and mutated residues are shown as blue and red colored bold letters, respectively.

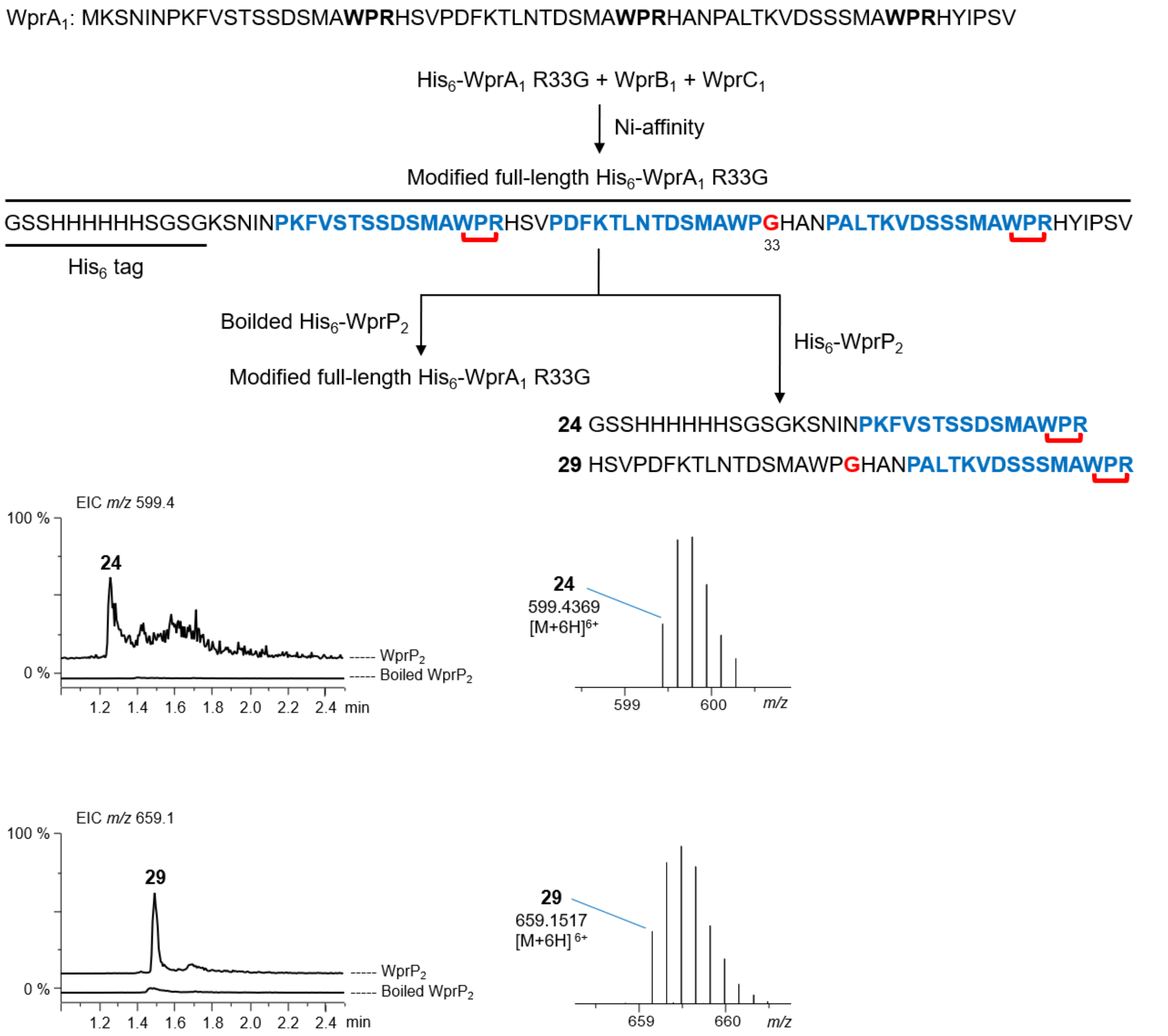


**Figure S25.** *In vitro* assay of His$_6$-WprP$_2$ or boiled His$_6$-WprP$_2$ + modified full-length His$_6$-WprA$_1$ R33G. The EIC chromatogram and MS spectra of fragments **24** and **29**. A single mutation at R33G on the precursor peptide prevented cleavage at this position and generated a longer fragment **29** containing two WPR motifs. Core peptides and mutated residues are shown as blue and red colored bold letters, respectively.

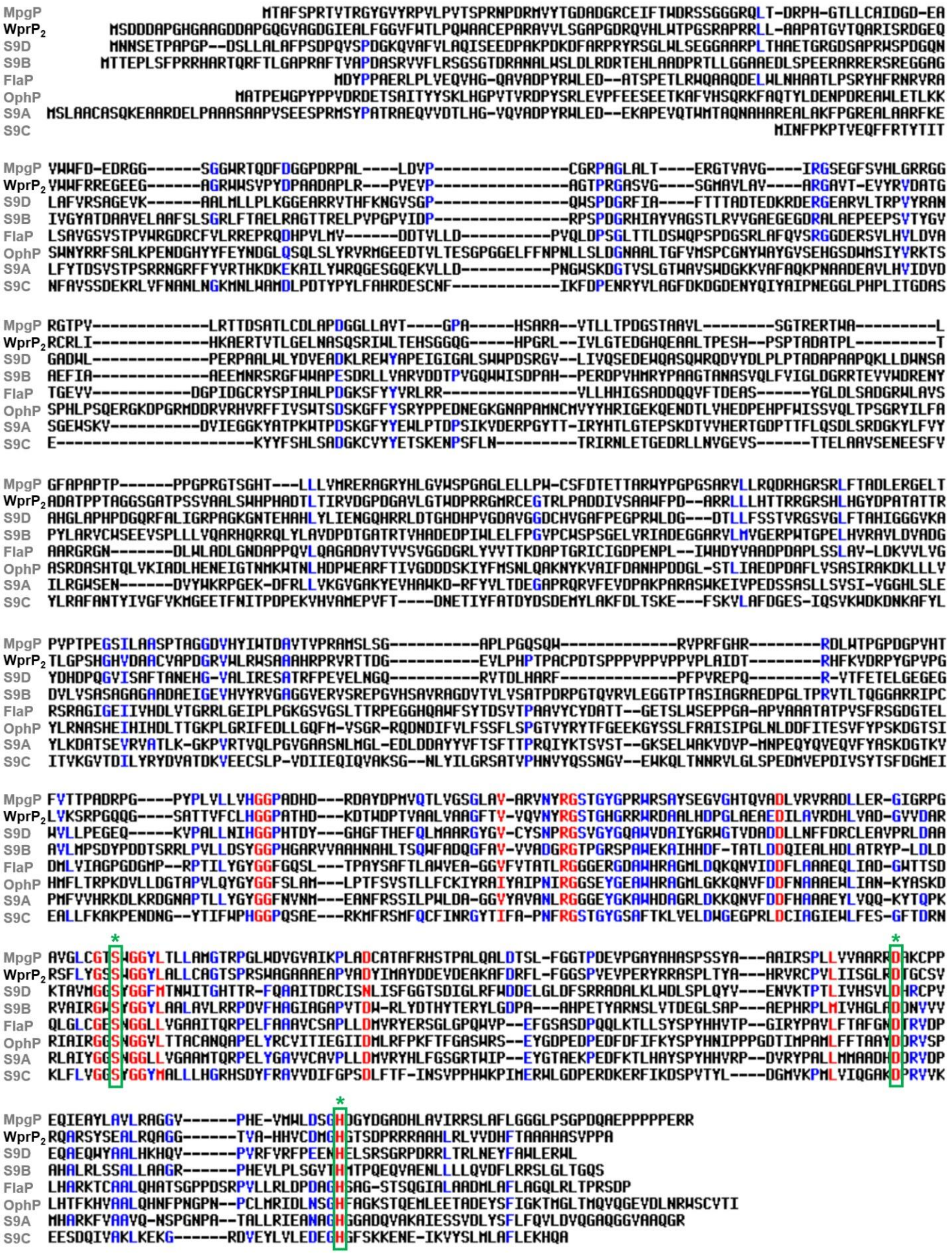


**Figure S26.** A sequence alignment generated using Multalin version 5.4.1.[21] Conserved catalytic triad consisting Ser, Asp, and His residues in known and newly identified S9 proteases are enclosed in green brackets. $WprP_2$ catalytic triad at Ser507, Asp590, and His621. Characterized S9 proteases FlaP (WP_012924050.1),[12] OphP (XPC16732.1),[13] and MpgP (WP_004993372.1).[14] The S9 subfamily proteases, S9A (WP_020478534.1; MEROPS ID S09.076), S9B (BAJ10544.1; MEROPS S09.073), S9C (WP_067208610.1), and S9D (WP_010886811.1; MEROPS ID S09.079) were obtained from the MEROPS database.[22]

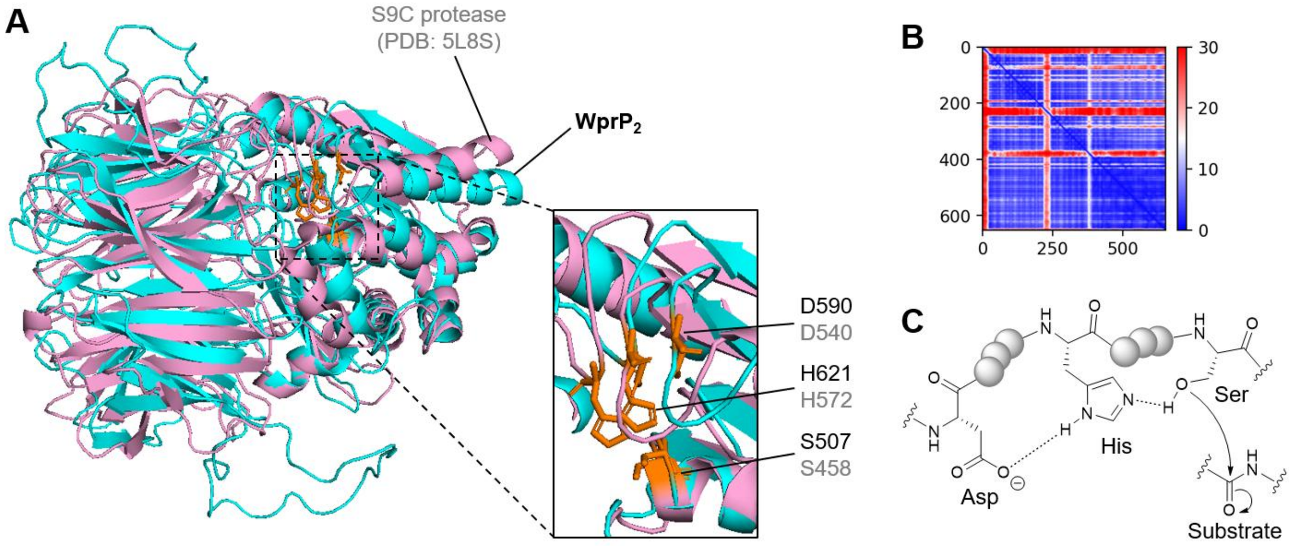


**Figure S27.** Overview of catalytic triad consisting of Ser, Asp and His residues in S9 proteases. (A) The crystal structure of C-terminal $His_6$ tag S9C acyl aminoacyl peptidase (PDB: 5L8S),[23] is superimposed with the predicted structure of S9 protease $WprP_2$ by ColabFold.[24] (B) AlphaFold2 confidence measure of PAE for modeling. (C) Typical mechanism of serine protease utilizing a catalytic triad (Ser, His, and Asp) to cleave peptide bonds of substrate.[25] Protein structures of 5L8S and $WprP_2$ are shown as pink and cyan colored ribbons, respectively.

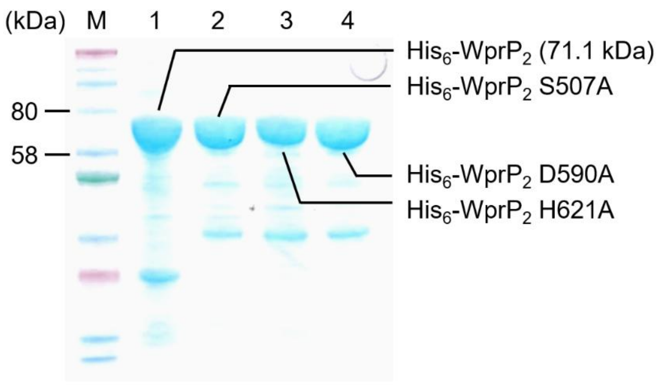


**Figure S28.** SDS-PAGE of $WprP_2$ mutants. M: molecular weight marker. 1: *in vivo* expression of $His_6$-$WprP_2$ in *E. coli* NiCo21 (DE3) and via Ni-affinity purification. 2: *in vivo* expression of $His_6$-$WprP_2$ S507A in *E. coli* NiCo21 (DE3) and via Ni-affinity purification. 3: *in vivo* expression of $His_6$-$WprP_2$ D590A in *E. coli* NiCo21 (DE3) and via Ni-affinity purification. 4: *in vivo* expression of $His_6$-$WprP_2$ H621A in *E. coli* NiCo21 (DE3) and via Ni-affinity purification. Theoretical molecular weight of recombinant $His_6$-$WprP_2$ is 71.1 kDa.

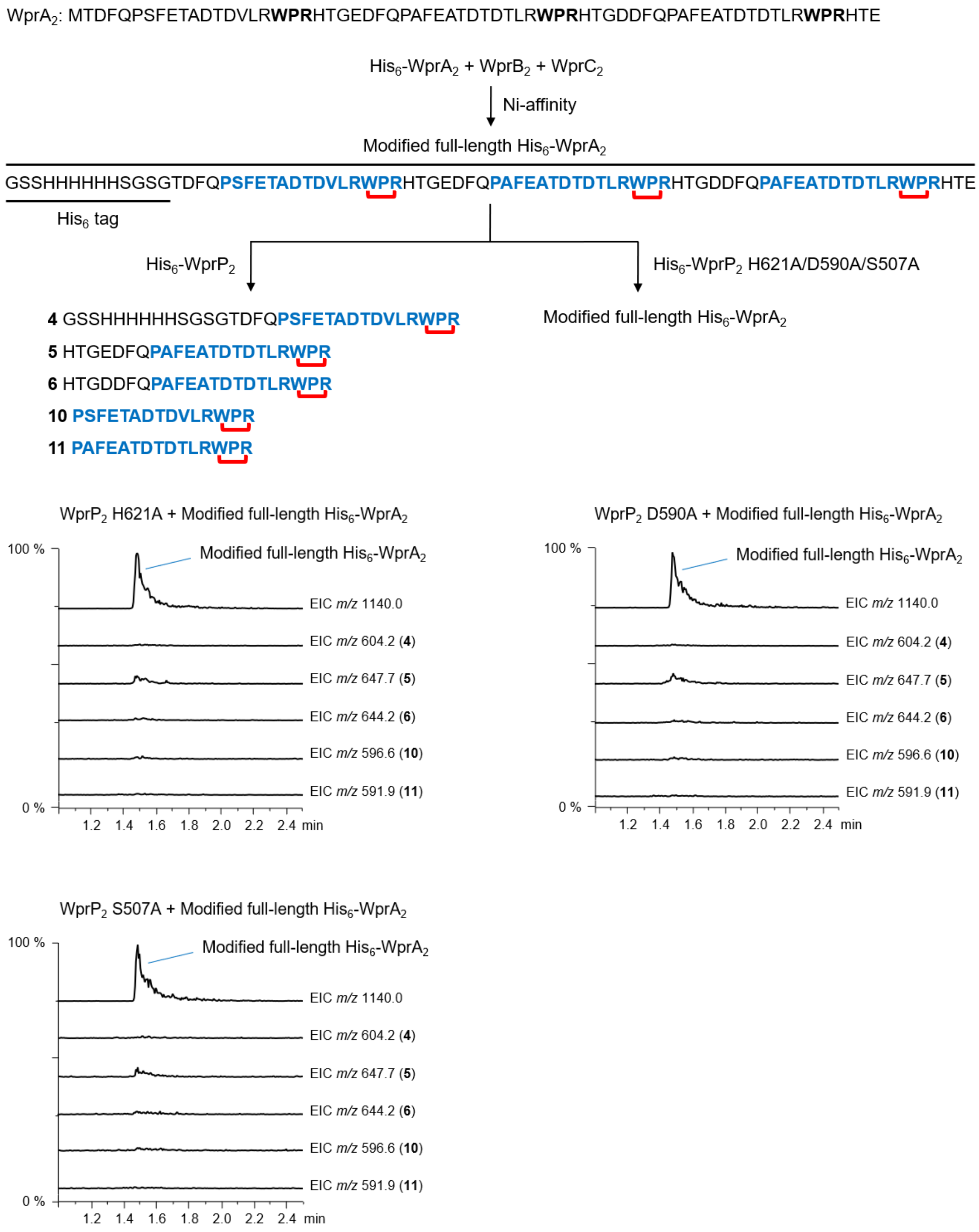


**Figure S29.** *In vitro* assay of His$_6$-WprP$_2$ H621A/D590A/S507A + modified full-length His$_6$-WprA$_1$. The EIC chromatogram of uncleaved full-length His$_6$-WprA$_1$ and all expected cleaved peptide fragments. A single mutation at H621A, D590A or S507A in $WprP_2$ abolished its cleavage activity. Core peptides are shown as blue colored bold letters.

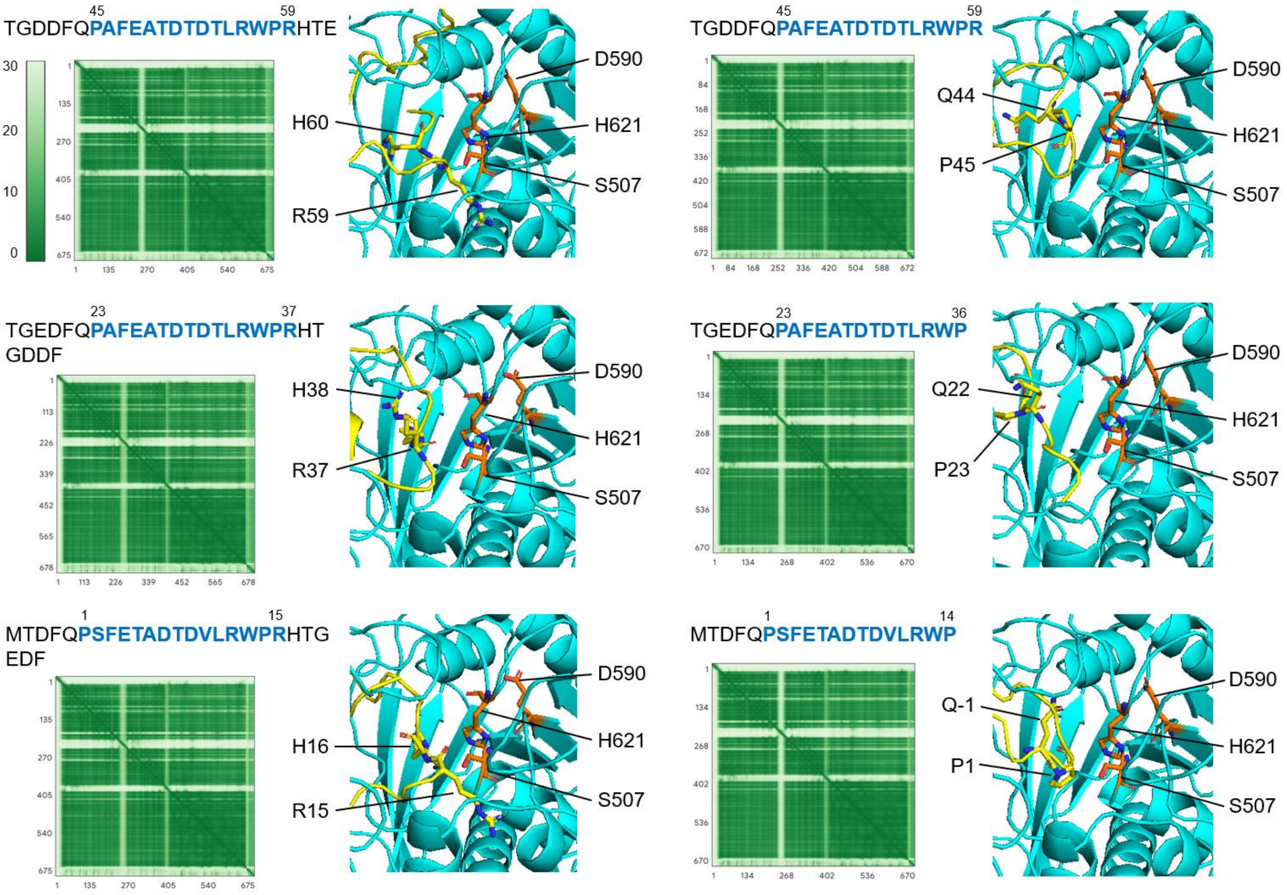


**Figure S30.** AlphaFold3 predicted the complex structure of $WprP_2$ with a truncated substrate of $WprA_2$.[26] The first cleavage sites of Arg59-His60, Arg37-His38 and Arg15-His16, and the second cleavage sites of Gln44-Pro45, Gln22-Pro23 and Gln-1-Pro1 are located in the catalytic triad. Structures of S9 protease $WprP_2$ and truncated substrate $WprA_2$ are shown as cyan and yellow colored ribbons, respectively.

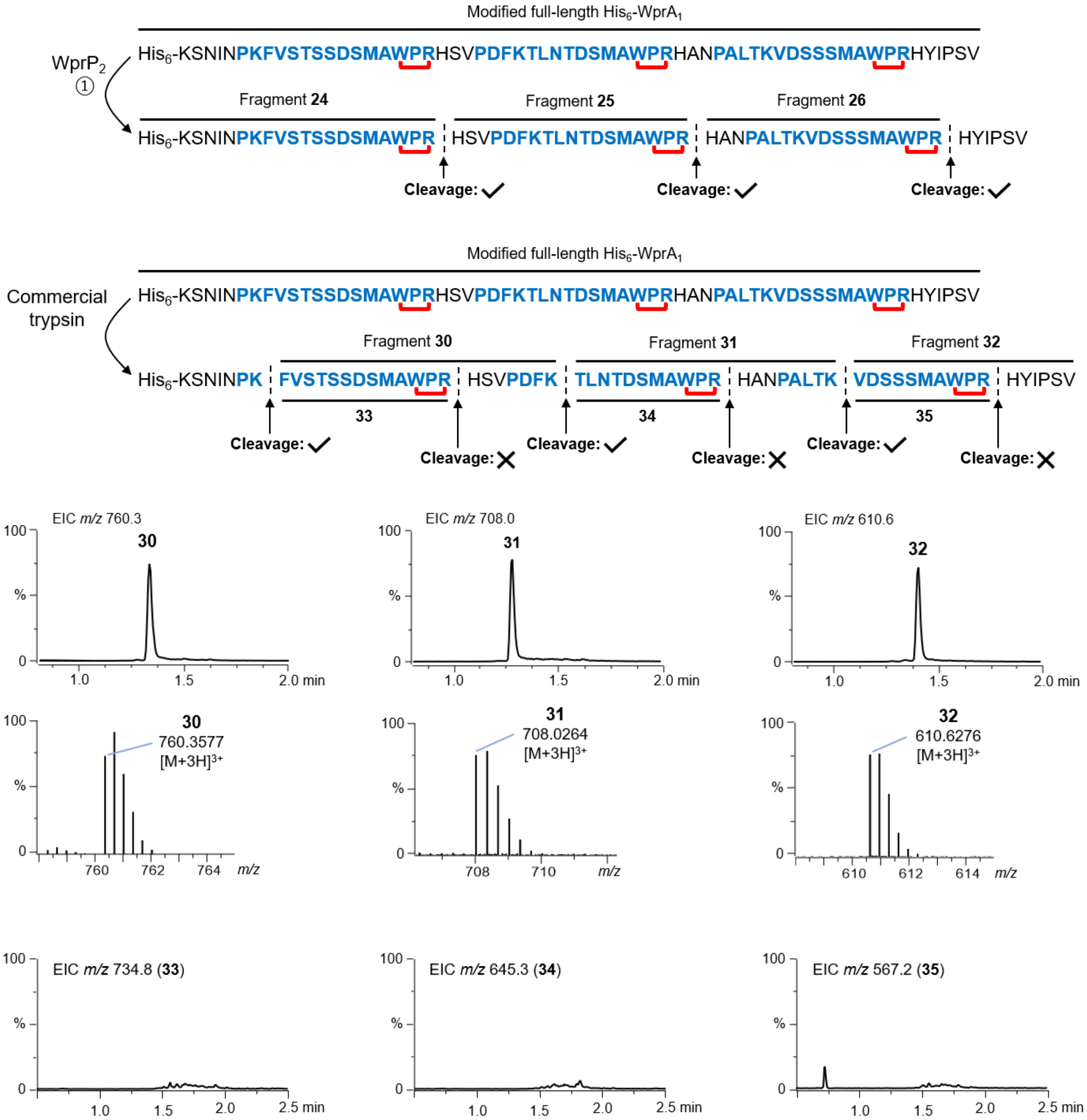


**Figure S31.** Overview of commercial trypsin and $WprP_2$ cleavage activity against modified full-length $His_6$-$WprA_1$. Trypsin is known to cleave after Arg or Lys residues.[27] Our previous data showed that trypsin can cleave the modified full-length $His_6$-$WprA_1$ at resides not involved in cross-link formation and generate fragments **30-32**.[2] Unlike $WprP_2$, commercial trypsin was unable to cleave the WPR-Xaa site of the modified full-length $WprA_1$, highlighting the potential applicability of $WprP_2$ in trypsin-like peptide bonds cleavage.

**Table S1**. List of 16 homologous WprB whose putative precursor peptides contain one or three repeated WPR motif.

| **Species** | **rSAM proteins** | **Precursor peptides** | **Putative S9 protease**[a] |
|---|---|---|---|
| *Xenorhabdus bovienii* | WP_274719294.1 | MKADFKPHFKMSNSESMA**WPR**HTVPS FETTNSQSMA**WPR**HATPSFTVADNTSM A**WPR**HKMSV | MDE1481349.1 MDE1481745.1 |
| *Xenorhabdus koppenhoeferi* | SFU55310.1 | MKNNLNPQFNRVGNDSMA**WPR**HVTVT PALQSVNSQSMA**WPR**HTTPALTKVENS AMA**WPR**HKMTD | SFU67181.1 SFU81005.1 |
| *Xenorhabdus koppenhoeferi* | WP_092550051.1 | MKNNLNPQFNRVGNDSMA**WPR**HVTVT PALQSVNSQSMA**WPR**HTTPALTKVENS AMA**WPR**HKMTD | WP_319940173.1 WP_422643826.1 |
| *Xenorhabdus* sp. | MDX7991759.1 | MKSNINPKFVSTSSDSMA**WPR**HSVPDF KTLNTDSMA**WPR**HANPALTKVDSSSMA **WPR**HYIPSV | MDX7992403.1 MDX7989948.1 |
| *Xenorhabdus littoralis* psl | WP_319939623.1 | MKSNINPKFVSTSSDSMA**WPR**HSVPDF KTLNTDSMA**WPR**HANPALTKVDSSSMA **WPR**HYIPSV | MDX7992403.1 MDX7989948.1 |
| *Xenorhabdus miraniensis* strain DSM 17902 | WP_099116060.1 | MKSILDPKFVNAGNDSMA**WPR**HSSPD FKVLNTESMA**WPR**HANPALTKAESSSM A**WPR**HYIPAN | WP_244170859.1 WP_422646015.1 |
| *Streptomyces jietaisiensis NBC_00023* | WP_331725114.1 | MVNRGLTDQHPGGGRVERVYRQVGR GNMETFAPSFEAPVGTDAMR**WPR**HTS | WTV91698.1 WTV90033.1 WTV91052.1 |
| *Streptomyces venezuelae* | WP_399537670.1 | MTDFQPSFETADTDVLR**WPR**HTGEDF QPAFEATDTDTLR**WPR**HTGDDFQPAFE ATDTDTLR**WPR**HTE | MFI7318140.1 MFI7323022.1 MFI7319834.1 MFI7322834.1 MFI7320687.1 |
| *Xenorhabdus bovienii* | WP_112238872.1 | MKADFKPHFKMSNSESMA**WPR**HTVPS FETTNSQSMA**WPR**HATPSFTVADNTSM A**WPR**HKMSV | |
| *Klebsiella pneumoniae* | HEF8906847.1 | MKTNMSPEFVIAGSESMA**WPR**HAVQDI IALNTDSMA**WPR**HAEPALTKAESSSMA **WPR**HYIPSV | HEF8905397.1 HEF8902083.1 HEF8904238.1 HEF8906957.1 |
| *Klebsiella quasipneumoniae* | HBR1386588.1 | MKTNMSPEFVIAGSESMA**WPR**HAVQDI IALNTDSMA**WPR**HAEPALTKAESSSMA **WPR**HYIPSV | HBR1384885.1 HBR1383797.1 |
| *Salmonella enterica* | EFP6323309.1 | MKTNISPEFVSAGSESMA**WPR**HAVSDL IALNTNSMA**WPR**HAEPALTKVESSSMA **WPR**HYIPSV | EFP6323746.1 EFP6319836.1 |
| *Photorhabdus heterorhabditis* | WP_149616604.1 | MKTNINPEFVNTRSDSMA**WPR**HAVPDI KALNTDSMA**WPR**HAVPALIKTESSSMA **WPR**HYIPAV | WP_149616992.1 WP_054481033.1 |
| *Klebsiella pneumoniae* | WP_394792610.1 | MKTNMSPEFVIAGSESMA**WPR**HAVQDI IALNTDSMA**WPR**HAEPALTKAESSSMA **WPR**HYIPSV | WP_394792317.1 WP_002895066.1 WP_224230348.1 |
| *Fastidiosibacter lacustris* strain NBRC 112274 | WP_116964676.1 | MKSSDNQKFAKASSDDPVV**YPR**HFAPD FKVMGAGSTQ**WPR**HANHVLTRSGVTS GCLSKT | n.d.[b] |
| *Vibrio* sp. | WP_341664272.1 | MKKLKVIKTTSSP**WPR**AKR | WP_341663497.1 WP_341661673.1 |

[a]The putative S9 protease was not found in the flanking regions of the homologous WprB enzyme but was found within the genome. [b]Not detected.

**Table S2**. Gene sequences used in this study.

| Gene | Vector (Restriction Sites) | Insert Sequence[b] |
|---|---|---|
| WprP$_2$[a] (WP_399537673.1) | pET28a(+) NcoI(-G)/XhoI | TCAGATGATGATGCTCCCGGACATGGCGCTGCTGGTGACG<br>ATGCTCCGGGTCAGGGTGTAGCGGGCGATGGCATAGAAG<br>CCCTGTTTGGCGGAGTTTTCTGGACGTTACCTCAGTGGGC<br>AGCGTGCGAACCCGCACGTGCAGTAGTGCTTTCAGGTGC<br>ACCTGGCGATCGGCAGGTGCACCTTTGGACACCTGGGAG<br>TCGTGCGCCACGTCGGTTACTTGCGGCTCCAGCCACTGG<br>AGTTACGCAGGCACGTATTAGCCGGGACGGAGAGCAGGT<br>CTGGTGGTTCCGCCGAGAGGGTGAAGAGGGAGCTGGAC<br>GATGGTGGTCAGTACCGTATGATCCGGCAGCTGACGCTCC<br>ACTTCGACCGGTAGAGGTTCCAGCGGGTACCCCGCGCGG<br>TGCAAGCGTTGGTAGCGGGATGGCGGTACTTGCAGTCGC<br>CCGCGGGGCAGTCACTGAAGTTTATCGGGTGGACGCCAC<br>AGGCAGATGCAGACTGATACATAAAGCTGAACGCACTGTC<br>ACACTCGGGGAACTGAATGCGTCCCAGAGCCGGATTTGG<br>CTTACAGAACATAGTGGTGGCCAAGGTCACCCAGGTCGCC<br>TGATTGTCTTAGGAACGGAAGATGGCCACCAGGAAGCGG<br>CGTTGACCCCAGAGAGCCACCCCAGTCCCACGGCGGATG<br>CTACACCATTGACCGCAGATGCGACCCCGCCAACAGCTG<br>GTGGATCAGGAGCAACCCCGAGCTCGGTTGCGGCGCTGA<br>GTTGGCATCCACATGCTGACACCTTAACCATCCGGGTTGA<br>TGGCCCAGATGGAGCCGTGCTTGGTACCTGGACCCACG<br>CCGCGGTATGCGCTGTGAAGGAACACGCCTTCCCGCCGA<br>TGATATAGTCAGCGCCGCCTGGTTTCCCGACGCTCGTCGT<br>TTATTGCTTCACACAACTCGCCGTGGTCGTTCACACTTACA<br>TGGATATGATCCCGCCACGGCGACAACACGTACTCTCGGA<br>CCGTCACACGGTCACGTTGATGCTGCGTGTGTCGCTCCT<br>GACGGACGAGTTTGGTTACGTTGGAGCGCGGCTGCCCAC<br>CGCCCACGGGTTCGAACTACAGATGGAGAAGTGTTACCAC<br>ATCCCACCCCCGCGTGTCCCGATACTTCTCCACCGCCTGT<br>GCCGCCTGTACCGCCTGTGCCGCTGGCCATCGATACACG<br>CCATTTTAAAGTTGACCGTCCATATGGTCCTGTTCCGGGTT<br>TAGTGAAGTCAAGACCTGGCCAGCAGGGTTCCGCAACGA<br>CGGTCTTTTGTTTACACGGTGGCCCGGCCACACATGATAA<br>AGACACTTGGGACCCTACCGTCGCAGCGCTTGTCGCCGC<br>GGGGTTTACTGTGGTTCAGGTCAATTATCGTGGCTCAACG<br>GGACATGGAAGACGGTGGCGCGATGCGGCCCTTCACGAT<br>CCGGGGCTCGCCGAGGCCGAGGATATCCTGGCGGTACGT<br>GACCATTTGGTTGCCGACGGGGTTGTTGATGCTCGAAGAT<br>CTTTCTTGTATGGTTCCTCATGGGGTGGATACCTTGCGCTT<br>TTGTGCGCTGGAACGTCACCCCGTTCCTGGGCGGGTGCT<br>GCAGCTGAAGCCCCGGTGGCCGACTATATAATGGCTTATG<br>ACGATGAAGTAGATGAGGCAAAAGCCTTTGACCGCTTTCT<br>GTTTGGCGGCTCGCCGGTAGAGGTGCCGGAACGGTATCG<br>TCGCGCTTCCCCCTTAACTTATGCGCACAGAGTCCGATGT<br>CCAGTCCTGATTATAAGTGGGTTACGTGATACCGGTTGCTC<br>GGTTCGTCAGGCACGTTCTTATTCGGAAGCTCTTAGACAG<br>GCAGGCGGCACGGTAGCGCATCATGTGTGTGATATGGGAC<br>ATGGAACCAGCGATCCCCGTCGTCGTGCGGCTCACCTGC<br>GCCTCGTGGTTGACCACTTTACGGCGGCAGCGCACGCCT<br>CTGTACCACCTGCCTAA |

| | | |
|---|---|---|
| WprB$_2$[a] (WP_399537670.1) | pCDFDuet-1 NdeI_XhoI | CAGACTGCGGGCATAAGACAGCAGCCGACTGGTGAACCT TTATTTGTTGGCCCGACTCATGTGGACCTTGACTTGACAAA CGCATGTAACTTGGCGTGTTCACATTGCTCCGTAGCGAGT GGAAAGCCCATGAAGGATGAGTTAGACACGGAAGGGATG CTGGGCGTTGTTCGCGACGTCCATGCCTTGGGTACGTTGT CATTGACGGTGGCGGGTGGAGAGCCTTTCATGCGAGAGG ACATTGTTGAACTGTTGGCTTTTGCTTGTTCCCTTCCTGGT TGGTCGGTAACAGTTATAACGAACGGAACATACTTCACAGA CCCGTTGCTTGACCAACTCCGTACTCGTTGTCCCGAGCTG ACCGTAAATGTCTCAGTAGACGGGAGCTCTCCGGAAACAT TCGACCGCCTGCGCCATCGCCGTAGACGTACACCTGAGG CGCAAAGACAATTGTTTGAACAAGTGACTCAAGGCATCCG GACAGCTGTTGACGCAGGCCTTAAGGTTCACACTTCATTC ACATTAGCGAGATGTAATGCAGACGATGTAGCGGCCACATA CGACCTCGTTCGAGGACTTGGGGCTAGAGACCTGCTGGC AATTAAGTTCTTCTCTGCGGGAAGAGGGTTGGATCACCTTA CTCAAATGGCGTTTCCGTACGCAGAGTGGGCTCGTACCAT GGTACGTTTGACCCGTCAGAAGGCAGCAGGACGTTTTCCT TTTCTTTCTCTGTCTTTACCTTCCGCTTGGGAATTCTATCTC CCCCTTCACGAGGCGGGAATGGACCTTAGACAAGCAGAG CGCTTATGGCGTTACCGTGCCCCGCTTCGCGGAAGCTATT ACGCTAGATTCCGCACAGTTGGGGACCCTTCAGGTGTGG CAGACCTTAACGTAGTTGCGAACGGCGACGTTTACCCTGT TACTCTGATGTCTGGTAACCGTGCGGCGCTTTGCGGGAAC GTGAGAACAACTCCGCTGGGCGACATATGGGAACACTCG CCCACGCTGGCAGCTCTGCGGGGATTGGACCTGAGAGAT CTTCCGCCAACCTGCGGTACATGCCCGGTAAGCACTCTGT GTGGTGGTGGCAGCCGTGCACGCGCCTTGATACGCAGTG GCAGCCTGGCTGGTCCGGATGCAAGTTGCCCAAAGCTCG ACACAGCCGAACGTACCAGCCACCAAAGTCACCGCTGA |
| WprA$_2$[a] (WP_399537669.1) | pACYCDuet-1 NcoI(-G)/ EcoRI | ggcagcagccatcaccatcatcaccacagcggcagcggcACGGACTTCCA ACCTAGCTTCGAGACGGCCGACACTGATGTGCTTAGATGG CCTCGCCATACAGGGGAAGATTTCCAGCCCGCCTTTGAAG CAACCGATACGGACACATTAAGATGGCCACGTCACACGGG TGACGACTTCCAGCCAGCGTTTGAGGCGACAGACACTGA CACACTGAGATGGCCGCGTCACACTGAGTGATCGAACAG AAAGTAATCGTATTGTACACGGCCGCATAATCGAAATTAATA CGACTCACTA |
| WprA$_2$C$_2$[a] (WP_399537669.1; WP_399537671.1) | pACYCDuet-1 NcoI(-G)/XhoI | ggcagcagccatcaccatcatcaccacagcggcagcggcACGGACTTCCA ACCTAGCTTCGAGACGGCCGACACTGATGTGCTTAGATGG CCTCGCCATACAGGGGAAGATTTCCAGCCCGCCTTTGAAG CAACCGATACGGACACATTAAGATGGCCACGTCACACGGG TGACGACTTCCAGCCAGCGTTTGAGGCGACAGACACTGA CACACTGAGATGGCCGCGTCACACTGAGTGATCGAACAG AAAGTAATCGTATTGTACACGGCCGCATAATCGAAATTAATA CGACTCACTATAGGGGAATTGTGAGCGGATAACAATTCCCC ATCTTAGTATATTAGTTAAGTATAAGAAGGAGATATACATatgT TAGTATTTGGCCGGGGAAGACACGCAGTTCGGGTATTGCC CAGACGAGAAGGTTGTGAAGTGCGTTTTGGCGAGTCAGTT GTCGTTGGAAACGAAGACCTTGCGGCTCTGCTGGCTTATG TACACTCCGATCCTACTGGGGAGCTTTTACCGCACGCAGC AGCTTTGCCAATGCCAGCGGAAGAATTCAGAACTTTAGCT CTTCGAACTCTGGCGAGCCTCGTCAGTGCTGGTGCGGAT |

| | | |
|---|---|---|
| | | CCGACAGTTTGGGGTCCTGGATGGACACCGCAAGCGGCG<br>GCGGATGGCCGAAGAGGAGCTTGA |
| WprA$_2$C$_2$-eng1[a] (D51A/T52A/D53A/T54A) | pACYCDuet-1 NcoI(-G)/XhoI | ggcagcagccatcaccatcatcaccacagcggcagcggcACGGACTTCCA<br>ACCTAGCTTCGAGACGGCCGACACTGATGTGCTTAGATGG<br>CCTCGCCATACAGGGGAAGATTTCCAGCCCGCCTTTGAAG<br>CAACCGATACGGACACATTAAGATGGCCACGTCACACGGG<br>TGACGACTTCCAGCCAGCGTTTGAGGCGACAgctgcagctgca<br>CTGAGATGGCCGCGTCACACTGAGTGATCGAACAGAAAGT<br>AATCGTATTGTACACGGCCGCATAATCGAAATTAATACGACT<br>CACTATAGGGGAATTGTGAGCGGATAACAATTCCCCATCTT<br>AGTATATTAGTTAAGTATAAGAAGGAGATATACATatgTTAGTA<br>TTTGGCCGGGGAAGACACGCAGTTCGGGTATTGCCCAGA<br>CGAGAAGGTTGTGAAGTGCGTTTTGGCGAGTCAGTTGTC<br>GTTGGAAACGAAGACCTTGCGGCTCTGCTGGCTTATGTAC<br>ACTCCGATCCTACTGGGGAGCTTTTACCGCACGCAGCAGC<br>TTTGCCAATGCCAGCGGAAGAATTCAGAACTTTAGCTCTTC<br>GAACTCTGGCGAGCCTCGTCAGTGCTGGTGCGGATCCGA<br>CAGTTTGGGGTCCTGGATGGACACCGCAAGCGGCGGCG<br>GATGGCCGAAGAGGAGCTTGA |
| WprA$_2$C$_2$-eng2[a] (D42A/F43A/P47A/E48A) | pACYCDuet-1 NcoI(-G)/XhoI | ggcagcagccatcaccatcatcaccacagcggcagcggcACGGACTTCCA<br>ACCTAGCTTCGAGACGGCCGACACTGATGTGCTTAGATGG<br>CCTCGCCATACAGGGGAAGATTTCCAGCCCGCCTTTGAAG<br>CAACCGATACGGACACATTAAGATGGCCACGTCACACGGG<br>TGACgcagctCAGCCAGCGgcagctGCGACAGACACTGACACA<br>CTGAGATGGCCGCGTCACACTGAGTGATCGAACAGAAAGT<br>AATCGTATTGTACACGGCCGCATAATCGAAATTAATACGACT<br>CACTATAGGGGAATTGTGAGCGGATAACAATTCCCCATCTT<br>AGTATATTAGTTAAGTATAAGAAGGAGATATACATatgTTAGTA<br>TTTGGCCGGGGAAGACACGCAGTTCGGGTATTGCCCAGA<br>CGAGAAGGTTGTGAAGTGCGTTTTGGCGAGTCAGTTGTC<br>GTTGGAAACGAAGACCTTGCGGCTCTGCTGGCTTATGTAC<br>ACTCCGATCCTACTGGGGAGCTTTTACCGCACGCAGCAGC<br>TTTGCCAATGCCAGCGGAAGAATTCAGAACTTTAGCTCTTC<br>GAACTCTGGCGAGCCTCGTCAGTGCTGGTGCGGATCCGA<br>CAGTTTGGGGTCCTGGATGGACACCGCAAGCGGCGGCG<br>GATGGCCGAAGAGGAGCTTGA |
| WprA$_2$C$_2$-eng3[a] (P45A) | pACYCDuet-1 NcoI(-G)/XhoI | ggcagcagccatcaccatcatcaccacagcggcagcggcACGGACTTCCA<br>ACCTAGCTTCGAGACGGCCGACACTGATGTGCTTAGATGG<br>CCTCGCCATACAGGGGAAGATTTCCAGCCCGCCTTTGAAG<br>CAACCGATACGGACACATTAAGATGGCCACGTCACACGGG<br>TGACGACTTCCAGgctGCGTTTGAGGCGACAGACACTGAC<br>ACACTGAGATGGCCGCGTCACACTGAGTGATCGAACAGAA<br>AGTAATCGTATTGTACACGGCCGCATAATCGAAATTAATACG<br>ACTCACTATAGGGGAATTGTGAGCGGATAACAATTCCCCAT<br>CTTAGTATATTAGTTAAGTATAAGAAGGAGATATACATatgTTA<br>GTATTTGGCCGGGGAAGACACGCAGTTCGGGTATTGCCCA<br>GACGAGAAGGTTGTGAAGTGCGTTTTGGCGAGTCAGTTGT<br>CGTTGGAAACGAAGACCTTGCGGCTCTGCTGGCTTATGTA<br>CACTCCGATCCTACTGGGGAGCTTTTACCGCACGCAGCAG<br>CTTTGCCAATGCCAGCGGAAGAATTCAGAACTTTAGCTCTT<br>CGAACTCTGGCGAGCCTCGTCAGTGCTGGTGCGGATCCG<br>ACAGTTTGGGGTCCTGGATGGACACCGCAAGCGGCGGCG<br>GATGGCCGAAGAGGAGCTTGA |

| | | |
|---|---|---|
| WprB1[a,c] (WP_3199396 23.1) | pCDFDuet-1 NdeI_XhoI | GAACATAATATTTCAGAGTGTATATCGCAGGACGAGCCCTT ATTCAATGCTCCAATACATGTGGACTTCGACATGACTAATG CATGCAATCTCGCATGTCACCACTGCCACGCCGCCAGTGG TAAGAGACAAAGAGACGAATTAACCTCGGAAGAGATCAAG AGAGCGATCGCAGAACTCCATCAAAACGGGGTAATCGACC TTACTATTGCTGGTGGGGAGCCCTTTCTGCGGCCGGAGTT GCCGGAAATCCTTGAGTACGCACACAGCTGTCAAGGAATT TATACCACCGTGGTGACAAATGGGACACTTTTAAAGCGCG ACACCGTAAAGAAATTGAGTTCTAACTGCAGCGGGGTTAA CTTTAATATAAGTATTGACGGCTCCACTCCGGATAAACTGG ATATTCTCCGTCACCGTAAGAAACGCGACAGTAATAAGCGC ACCCTGCTGTTCCGTCAAGTGACTGCAGGTGCCAAGGCA ATTAGCGACGCGGGACTTACTCTCGGAGTTAGCTTTGTGG TGTCTGGTATGAACGCCGACGACTTAGAAAATGTATACGAT ATGGCTGTTAACGAGTTGGGAGCCAAATCAGTTACCGCGA TCCGCTTCTTCCCAGCGGGTTTCGGTAAGTCTGCTCTGTC AGAATTAGCGCTCAAGTACGAGGAGTGGGAGAAAATCATC TTAGACCTGACAAGACGTCATGACTCTTTCCAAAACTTAAC CATTTCCGTCTCTGCCCCATGGGAGATTTATCTGCCGTTGC TTAACAATGGATATACACCGCAGCAAGTGTGGGAAATTTGG CAGTATCGCAGCACTCTTACCGACCCCGTGTACTCTTCGG AGTACCAAGTGGGCGATGCCAGCGGGATAGGTGATCTGAA CATCTCAGGTAACGGTAAGGTTTACCCATCGGTTCTTATGA GCGGTAATCACGACGTGCTGTGTGGCAACATACGGGAACA GAGTCTCAAGGAAATTTGGTACAACTCATATACACTTAAGA AACTCCGAAATATACGGCTTTCAGAAATTGGTGGTCCGTGT TTAGACTGTCATATACGTGACTTGTGCGGGGCAGGCTCTC GATCAAGAGCACTGTCCATTACAGGCTCCATCTACGGCCT CGACCACTGGTGTCCGGTTATTGCCGAAACCTACAGACGG AAGAATCACGTCATCGCCTTGCAAAAGCAATAA |
| WprA1C1[a,c] (Not annotated by NCBI; WP_31993962 2.1) | pACYCDuet-1 NcoI(-G)/XhoI | ggcagcagccatcaccatcatcaccacagcggcagcggcAAGTCTAACATC AACCCAAAGTTCGTTTCGACGTCAAGTGACTCAATGGCGT GGCCTCGTCATTCCGTACCGGACTTCAAGACATTAAACAC GGACTCAATGGCCTGGCCCCGTCATGCCAACCCCGCACT TACGAAAGTGGACTCATCATCGATGGCGTGGCCGCGCCAT TACATCCCATCAGTGTGATCGAACAGAAAGTAATCGTATTG TACACGGCCGCATAATCGAAATTAATACGACTCACTATAGG GGAATTGTGAGCGGATAACAATTCCCCATCTTAGTATATTAG TTAAGTATAAGAAGGAGATATACATATGTTATTGCACTCGCG AAACAATAACACCTTACGTTTACTCGAGCGACACTACGGG GCGGAGCTGCGTTACAATAGACAGGTCTTCTTCGCTAACA ATGACTTGGCCCGTATCATTAAGTTCGTCATCGAGACAAGC GATTGTGACTACCTTCCGCTGTGCCAACAGGGCCTGGAC GAGAAGATTGTTAGAAACACGCTGCACAAGCTCTCCGAGG TTGGTATCCCATTGTCGTTTTGGGGAGCGATATGGTCAGAC ACAGAAGACTCAAGCGCAAAGCGTGACGGGCTCTGA |
| WprA1C1-eng2[a] (H52A) (Not annotated by NCBI; WP_31993962 2.1) | pACYCDuet-1 NcoI(-G)/XhoI | ggcagcagccatcaccatcatcaccacagcggcagcggcAAGTCTAACATC AACCCAAAGTTCGTTTCGACGTCAAGTGACTCAATGGCGT GGCCTCGTCATTCCGTACCGGACTTCAAGACATTAAACAC GGACTCAATGGCCTGGCCCCGTCATGCCAACCCCGCACT TACGAAAGTGGACTCATCATCGATGGCGTGGCCGCGCgctT ACATCCCATCAGTGTGATCGAACAGAAAGTAATCGTATTGT ACACGGCCGCATAATCGAAATTAATACGACTCACTATAGGG |

| | | |
|---|---|---|
| | | GAATTGTGAGCGGATAACAATTCCCCATCTTAGTATATTAGTTAAGTATAAGAAGGAGATATACATATGTTATTGCACTCGCGAAACAATAACACCTTACGTTTACTCGAGCGACACTACGGGGCGGAGCTGCGTTACAATAGACAGGTCTTCTTCGCTAACAATGACTTGGCCCGTATCATTAAGTTCGTCATCGAGACAAGCGATTGTGACTACCTTCCGCTGTGCCAACAGGGCCTGGACGAGAAGATTGTTAGAAACACGCTGCACAAGCTCTCCGAGGTTGGTATCCCATTGTCGTTTTGGGGAGCGATATGGTCAGACACAGAAGACTCAAGCGCAAAGCGTGACGGGCTCTGA |

[a]Codons were optimized for heterologous expression in *E. coli*. [b]$His_6$ tag sequence was described in blue colored letters. [c]Plasmids from our previous study.[2]

**Table S3**. Amino acid sequence of proteins used in this study.

| **Protein** | **sequence** |
|---|---|
| $WprP_2$ | MSDDDAPGHGAAGDDAPGQGVAGDGIEALFGGVFWTLPQWAACEPARAVVLSGAPGDRQVHLWTPGSRAPRRLLAAPATGVTQARISRDGEQVWWFRREGEEGAGRWWSVPYDPAADAPLRPVEVPAGTPRGASVGSGMAVLAVARGAVTEVYRVDATGRCRLIHKAERTVTLGELNASQSRIWLTEHSGGQGHPGRLIVLGTEDGHQEAALTPESHPSPTADATPLTADATPPTAGGSGATPSSVAALSWHPHADTLTIRVDGPDGAVLGTWDPRRGMRCEGTRLPADDIVSAAWFPDARRLLLHTTRRGRSHLHGYDPATATTRTLGPSHGHVDAACVAPDGRVWLRWSAAAHRPRVRTTDGEVLPHPTPACPDTSPPPVPPVPPVPLAIDTRHFKVDRPYGPVPGLVKSRPGQQGSATTVFCLHGGPATHDKDTWDPTVAALVAAGFTVVQVNYRGSTGHGRRWRDAALHDPGLAEAEDILAVRDHLVADGVVDARRSFLYGSSWGGYLALLCAGTSPRSWAGAAAEAPVADYIMAYDDEVDEAKAFDRFLFGGSPVEVPERYRRASPLTYAHRVRCPVLIISGLRDTGCSVRQARSYSEALRQAGGTVAHHVCDMGHGTSDPRRRAAHLRLVVDHFTAAAHASVPPA |
| $WprB_2$ | MQTAGIRQQPTGEPLFVGPTHVDLDLTNACNLACSHCSVASGKPMKDELDTEGMLGVVRDVHALGTLSLTVAGGEPFMREDIVELLAFACSLPGWSVTVITNGTYFTDPLLDQLRTRCPELTVNVSVDGSSPETFDRLRHRRRRTPEAQRQLFEQVTQGIRTAVDAGLKVHTSFTLARCNADDVAATYDLVRGLGARDLLAIKFFSAGRGLDHLTQMAFPYAEWARTMVRLTRQKAAGRFPFLSLSLPSAWEFYLPLHEAGMDLRQAERLWRYRAPLRGSYYARFRTVGDPSGVADLNVVANGDVYPVTLMSGNRAALCGNVRTTPLGDIWEHSPTLAALRGLDLRDLPPTCGTCPVSTLCGGGSRARALIRSGSLAGPDASCPKLDTAERTSHQSHR |
| $WprB_1$ | MEHNISECISQDEPLFNAPIHVDFDMTNACNLACHHCHAASGKRQRDELTSEEIKRAIAELHQNGVIDLTIAGGEPFLRPELPEILEYAHSCQGIYTTVVTNGTLLKRDTVKKLSSNCSGVNFNISIDGSTPDKLDILRHRKKRDSNKRTLLFRQVTAGAKAISDAGLTLGVSFVVSGMNADDLENVYDMAVNELGAKSVTAIRFFPAGFGKSALSELALKYEEWEKIILDLTRRHDSFQNLTISVSAPWEIYLPLLNNGYTPQQVWEIWQYRSTLTDPVYSSEYQVGDASGIGDLNISGNGKVYPSVLMSGNHDVLCGNIREQSLKEIWYNSYTLKKLRNIRLSEIGGPCLDCHIRDLCGAGSRSRALSITGSIYGLDHWCPVIAETYRRKNHVIALQKQ |

**Table S4**. Primers used in this study.

| Primer | Sequence |
|---|---|
| $WprA_1$ R33G Fw[a] | CATGCCAACCCCGCACTTAC |
| $WprA_1$ R33G Rv[a] | accGGGCCAGGCCATTGAGTC |
| $WprA_1$ N36Q Fw | cagcccgcacttacgaaagtggac |
| $WprA_1$ N36Q Rv | ggcatgacggggccaggc |
| $WprA_2$ Q44A Fw | gcgccagcgtttgaggcg |
| $WprA_2$ Q44A Rv | gaagtcgtcacccgtgtgacg |
| $WprA_2$ H38A Fw | gccacgggtgacgacttcc |
| $WprA_2$ H38A Rv | acgtggccatcttaatgtgtccg |
| $WprA_2$ R59A Fw | gctCACACTGAGTGATCGAACAG |
| $WprA_2$ R59A Rv | CGGCCATCTCAGTGTGTC |
| $WprP_2$ H621A Fw | gctGGAACCAGCGATCCCCGT |
| $WprP_2$ H621A Rv | TCCCATATCACACACATGATGCG |
| $WprP_2$ D590A Fw | gctACCGGTTGCTCGGTTCGT |
| $WprP_2$ D590A Rv | ACGTAACCCACTTATAATCAGGACTG |
| $WprP_2$ S507A Fw | gctTGGGGTGGATACCTTGCG |
| $WprP_2$ S507A Rv | GGAACCATACAAGAAAGATCTTCG |

[a]Plasmid constructed from our previous study.[2]

**Table S5**. Plasmids constructed in this study.

| Plasmids | Description |
|---|---|
| $WprA_1$ (R33G) $C_1$-pACYCDuet-1[a] | Plasmid for expression of $WprA_1$ R33G + $WprC_1$ |
| $WprA_1$ (H52A) $C_1$-pACYCDuet-1 | Plasmid for expression of $WprA_1$ H52A + $WprC_1$ |
| $WprA_2$ (Q44A) $C_1$-pACYCDuet-1 | Plasmid for expression of $WprA_2$ Q44A + $WprC_2$ |
| $WprA_2$ (H38A) $C_1$-pACYCDuet-1 | Plasmid for expression of $WprA_2$ H38A + $WprC_2$ |
| $WprA_2$ (R59A) $C_1$-pACYCDuet-1 | Plasmid for expression of $WprA_2$ R59A + $WprC_2$ |
| $WprP_2$ H621A-pET28a(+) | Plasmid for expression of $WprP_2$ H621A |
| $WprP_2$ D590A-pET28a(+) | Plasmid for expression of $WprP_2$ D590A |
| $WprP_2$ S507A-pET28a(+) | Plasmid for expression of $WprP_2$ S507A |

[c]Plasmid constructed from our previous study.[2]

**Table S6**. Strains used in this study.

| Strains | Description |
|---|---|
| *E. coli* $WprA_2$ | *E. coli* NiCo (DE3) harboring $WprA_2$-pACYCDuet-1 |
| *E. coli* $WprA_2$ + $WprB_2$ | *E. coli* NiCo (DE3) harboring $WprA_2$-pACYCDuet-1 and $WprB_2$-pCDFDuet-1 |
| *E. coli* $WprA_2$ + $WprB_2$ + $WprC_2$ | *E. coli* NiCo (DE3) harboring $WprA_2C_2$-pACYCDuet-1 and $WprB_2$-pCDFDuet-1 |
| *E. coli* $WprA_2$-eng1 + $WprB_2$ + $WprC_2$ | *E. coli* NiCo (DE3) harboring $WprA_2$ (D51A/T52A/D53A/T54A) $C_2$-pACYCDuet-1 and $WprB_2$-pCDFDuet-1 |
| *E. coli* $WprA_2$-eng2 + $WprB_2$ + $WprC_2$ | *E. coli* NiCo (DE3) harboring $WprA_2$ (D42A/F43A/P47A/E48A) $C_2$-pACYCDuet-1 and $WprB_2$-pCDFDuet-1 |
| *E. coli* $WprA_2$ P45A + $WprB_2$ + $WprC_2$ | *E. coli* NiCo (DE3) harboring $WprA_2$ (P45A) $C_2$-pACYCDuet-1 and $WprB_2$-pCDFDuet-1 |
| *E. coli* $WprA_2$ Q44A + $WprB_2$ + $WprC_2$ | *E. coli* NiCo (DE3) harboring $WprA_2$ (Q44A) $C_2$-pACYCDuet-1 and $WprB_2$-pCDFDuet-1 |
| *E. coli* $WprA_2$ H38A + $WprB_2$ + $WprC_2$ | *E. coli* NiCo (DE3) harboring $WprA_2$ (H38A) $C_2$-pACYCDuet-1 and $WprB_2$-pCDFDuet-1 |
| *E. coli* $WprA_2$ R59A + $WprB_2$ + $WprC_2$ | *E. coli* NiCo (DE3) harboring $WprA_2$ (R59A) $C_2$-pACYCDuet-1 and $WprB_2$-pCDFDuet-1 |
| *E. coli* $WprA_1$ + $WprB_1$ + $WprC_1$[a] | *E. coli* NiCo (DE3) harboring $WprA_1C_1$-pACYCDuet-1 and $WprB_1$-pCDFDuet-1 |
| *E. coli* $WprA_1$ R33G + $WprB_1$ + $WprC_1$[a] | *E. coli* NiCo (DE3) harboring $WprA_1$ (R33G) $C_1$-pACYCDuet-1 and $WprB_1$-pCDFDuet-1 |
| *E. coli* $WprA_1$ H52A + $WprB_1$ + $WprC_1$ | *E. coli* NiCo (DE3) harboring $WprA_1$ (H52A) $C_1$-pACYCDuet-1 and $WprB_1$-pCDFDuet-1 |
| *E. coli* $WprA_1$ N36Q + $WprB_1$ + $WprC_1$ | *E. coli* NiCo (DE3) harboring $WprA_1$ (N36Q) $C_1$-pACYCDuet-1 and $WprB_1$-pCDFDuet-1 |
| *E. coli* $WprP_2$ | *E. coli* NiCo (DE3) harboring $WprP_2$-pET28a(+) |
| *E. coli* $WprP_2$ H621A | *E. coli* NiCo (DE3) harboring $WprP_2$ H621A-pET28a(+) |
| *E. coli* $WprP_2$ D590A | *E. coli* NiCo (DE3) harboring $WprP_2$ D590A-pET28a(+) |
| *E. coli* $WprP_2$ S507A | *E. coli* NiCo (DE3) harboring $WprP_2$ S507A-pET28a(+) |

[a]Strains from our previous study.[2]

**Table S7**. Precursor peptides used in this study. Red color residues indicate mutated residues.

| **Precursor Peptides** | **Amino Acids Sequence** |
|---|---|
| WprA$_2$ | MTDFQPSFETADTDVLR**WPR**HTGEDFQPAFEATDTDTLR**WPR** HTGDDFQPAFEATDTDTLR**WPR**HTE |
| WprA$_2$-eng1 (D51A/T52A/D53A/T54A) | MTDFQPSFETADTDVLR**WPR**HTGEDFQPAFEATDTDTLR**WPR** HTGDDFQPAFEATAAAALR**WPR**HTE |
| WprA$_2$-eng2 (D42A/F43A/P47A/E48A) | MTDFQPSFETADTDVLR**WPR**HTGEDFQPAFEATDTDTLR**WPR** HTGDAAQPAAAATDTDTLR**WPR**HTE |
| WprA$_2$ P45A | MTDFQPSFETADTDVLR**WPR**HTGEDFQPAFEATDTDTLR**WPR** HTGDDFQAAFEATDTDTLR**WPR**HTE |
| WprA$_2$ Q44A | MTDFQPSFETADTDVLR**WPR**HTGEDFQPAFEATDTDTLR**WPR** HTGDDF**A**PAFEATDTDTLR**WPR**HTE |
| WprA$_2$ H38A | MTDFQPSFETADTDVLR**WPR**HTGEDFQPAFEATDTDTLR**WPR** **A**TGDDFQPAFEATDTDTLR**WPR**HTE |
| WprA$_2$ R59A | MTDFQPSFETADTDVLR**WPR**HTGEDFQPAFEATDTDTLR**WPR** HTGDDFQPAFEATDTDTLR**WPA**HTE |
| WprA$_1$[a] | MKSNINPKFVSTSSDSMA**WPR**HSVPDFKTLNTDSMA**WPR**HAN PALTKVDSSSMA**WPR**HYIPSV |
| WprA$_1$ R33G | MKSNINPKFVSTSSDSMA**WPR**HSVPDFKTLNTDSMA**WPG**HAN PALTKVDSSSMA**WPR**HYIPSV |
| WprA$_1$ H52A | MKSNINPKFVSTSSDSMA**WPR**HSVPDFKTLNTDSMA**WPR**HAN PALTKVDSSSMA**WPRA**YIPSV |
| WprA$_1$ N36Q | MKSNINPKFVSTSSDSMA**WPR**HSVPDFKTLNTDSMA**WPR**HA**Q** PALTKVDSSSMA**WPR**HYIPSV |

[a]Precursor peptides from our previous study.[2]
